# Synthetic Modular Cyanobacterial Consortium Enables Carbon-Negative Production of Glycerol and Derivatives

**DOI:** 10.64898/2025.12.04.692262

**Authors:** Shubin Li, Tao Sun, Dailin Liu, Kungang Pan, Chen Lei, Weiwen Zhang

## Abstract

Cyanobacteria offer a direct route for converting solar energy and CO_2_ into valuable chemicals, yet the metabolic burden of complex heterologous pathways often limits their productivity and stability. Here, we present a modular co-culture strategy that uses glycerol as an efficient mediator metabolite to link cyanobacterial carbon fixation with downstream bioconversion. We first engineered *Synechococcus elongatus* UTEX 2973 (Syn2973) for high-level glycerol biosynthesis by optimizing synthase selection, eliminating competing pathways, and improving CO_2_-supply conditions, achieving a production of 14.53 g·L^-1^. In parallel, we constructed glycerol conversion modules in Syn2973 and *Gluconobacter oxydans* enabling production of 1,3-propanediol (1,3-PDO), 3-hydroxypropionic acid, and dihydroxyacetone, with exogenous glycerol yielding up to 30.07 g·L^-1^ 1,3-PDO. Integrating the two modules resulted in a fully autotrophic co-culture that produced all three target chemicals directly from CO_2_, including semi-continuous 1,3-PDO production at 6.02 g·L^-1^·day^-1^, corresponding to a net carbon fixation efficiency of 3.43 kg CO_2_ eq/kg 1,3-PDO.

## Introduction

In 2024, global energy-related CO_2_ emissions increased by 0.9% compared to 2023, reaching a record-high of 36.3 Gt ^1^. At this pace, projections suggest that the threshold for limiting warming to 1.5 °C could be breached within the next five years ^1^. Achieving carbon neutrality ahead of schedule has therefore become a global imperative ^2^. Photosynthesis, the largest energy and matter transformation process on Earth, assimilates over 300 Gt of CO_2_ annually ^3^. Cyanobacteria, the only prokaryotes capable of oxygenic photosynthesis, account for about 25% of this global carbon fixation ^4–6^. Model strains such as *Synechocystis* sp. PCC 6803 and *Synechococcus elongatus* PCC 7942 possess compact genomes and are highly tractable, enabling their use as chassis for synthetic biology. Through cell engineering, these organisms can be rewired to directly convert solar energy and CO_2_ into biofuels and high-value chemicals^7–10^, positioning them as promising platforms for carbon-negative biomanufacturing.

*Synechococcus elongatus* UTEX 2973 (hereafter Syn2973) is a fast-growing cyanobacterium recently identified as a promising microbial chassis. Under optimized conditions, Syn2973 exhibits a doubling time as short as 1.5 hours, and demonstrates remarkable tolerance to high temperature (up to 45 °C) and high light intensity (2000 μmol photons m^-2^ s^-1^) ^11^. Biomass concentration up to 23.41 g·L^-1^ have been achieved under semi-continuous cultivation ^12^. These traits highlight the potential of Syn2973 as an efficient photoautotrophic platform for sustainable bio-manufacturing. However, compared to *Escherichia coli* or *Saccharomyces cerevisiae*, cyanobacteria remain challenging to genetically engineer^13, 14^. Complex heterologous pathways often impose heavy metabolic burdens or generate toxic effects, leading to genetic instability and limited scalability for the engineered strains. To address these limitations, modular co-culture strategies have gained traction. By distributing pathway segments across distinct microbial partners, co-culture systems can reduce intracellular burden, alleviate regulatory interference, and provide flexible pathway design ^15, 16^.

Syn2973 is highly efficient at fixing CO_2_ into organic carbon, and its engineered sucrose production can reach titers of ∼8 g·L^-1^ ^17^. Building upon this, numerous studies have explored the use of sucrose as a mediator metabolite, coupling a cyanobacterial carbon fixation module with a heterotrophic conversion module to achieve CO_2_-to-chemical transformation. For instance, Li *et al.* demonstrated the co-culture of sucrose-producing *S. elongatus* PCC 7942 with *Vibrio natriegens* to produce lactic acid, *p*-coumaric acid, and 2,3-butanediol ^18^. However, few heterotrophic microbes can naturally utilize sucrose, and artificially introduced sucrose utilization pathways often exhibit poor efficiency. Therefore, the identification of more suitable mediator metabolites remains a critical challenge. Notably, cinnamic acid has also been used as a shuttle compound for downstream synthesis of styrene and other products ^19^, although its biosynthesis in cyanobacteria remains limited.

In this study (**Fig. 1**), we investigated glycerol as an alternative and advantageous mediator metabolite. Glycerol is readily metabolized by diverse microorganisms and serves as a key intermediate for the synthesis of high-value products such as 1,3-propanediol (1,3-PDO) and 3-hydroxypropionic acid (3-HP) ^20^. Importantly, its precursor glyceraldehyde-3-phosphate is a direct product of the Calvin-Benson-Bassham (CBB) cycle, suggesting strong potential for high-yield production in photosynthetic hosts. Based on this rationale, we constructed a two-module system comprising a glycerol production module and a glycerol conversion module. In the glycerol production module, Syn2973 was engineered for enhanced glycerol biosynthesis by optimizing synthase selection, blocking competing pathways, and improving CO_2_ supply strategies, achieving a glycerol production of 14.53 g·L^-1^. In the glycerol conversion module, we established three downstream pathways for the production of 1,3-propanediol (1,3-PDO), 3-hydroxypropionic acid (3-HP), and dihydroxyacetone (DHA), using either engineered Syn2973 or *Gluconobacter oxydans*. Via systematic metabolic engineering, the maximal production upon exogenous glycerol supplementation reached 30.07 g·L^-1^ for 1,3-PDO. Finally, the two modules were integrated into a co-culture system using CO_2_ as the sole carbon source. The synthetic modular glycerol-mediated system achieved *de novo* production of 1,3-PDO, 3-HP, and DHA. Meanwhile, under semi-continuous co-culture, 1,3-propanediol was stably produced stably at 6.02 g·L^-1^·day^-1^. Overall, this study establishes a modular, glycerol-mediated framework that significantly expands the capability of cyanobacterial platforms for sustainable biomanufacturing.

**Fig. 1.**
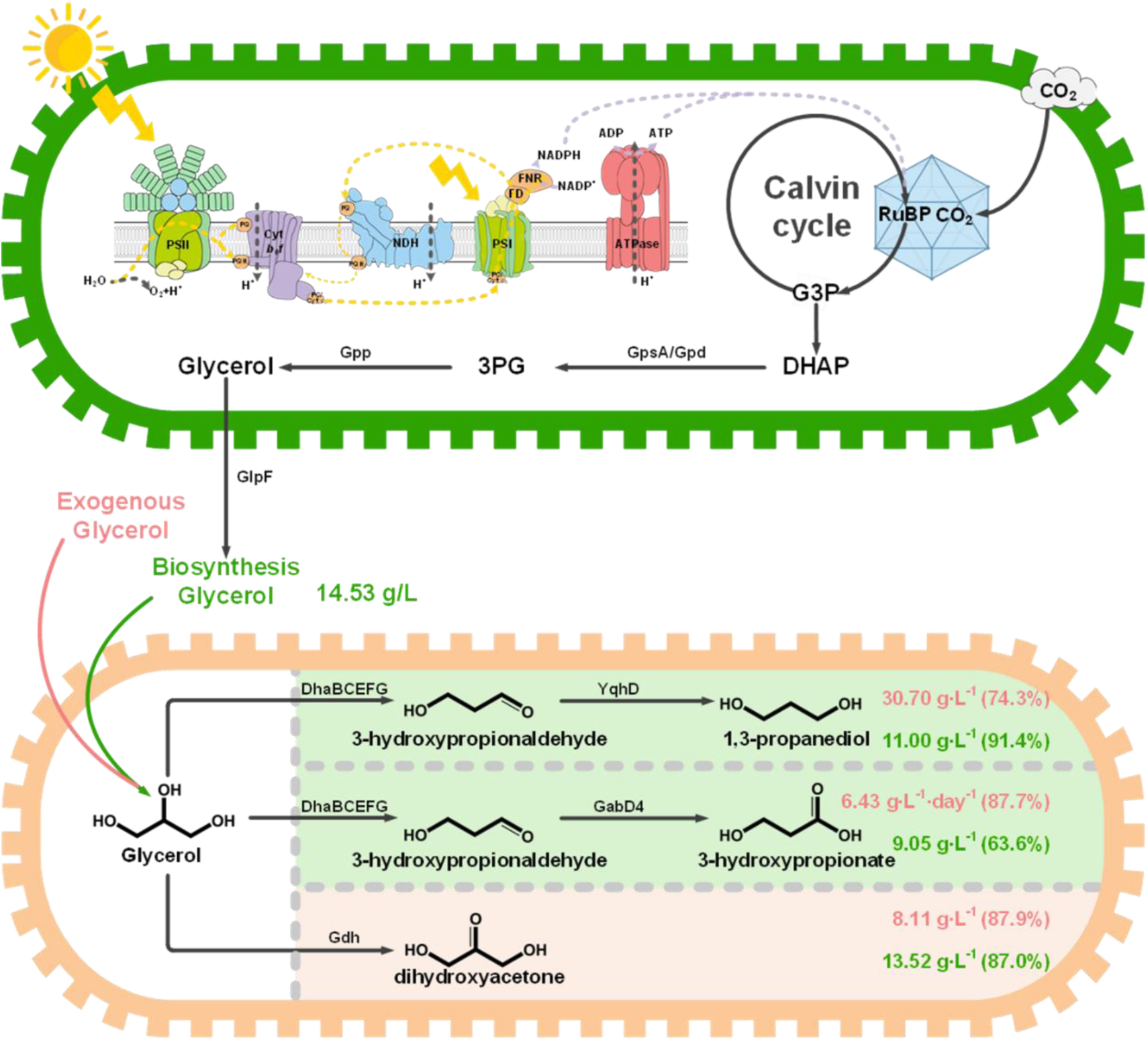
Schematic of this study.

## Results

### Pathway engineering of Syn2973 achieves 14.53 g·L^-1^ glycerol production

As shown in **Fig. 2A**, glyceraldehyde-3-phosphate (G3P), an intermediate of the CBB cycle, can be converted to glycerol through a three-step pathway involving triosephosphate isomerase (Tpi), glycerol-3-phosphate dehydrogenase (Gpd), and glycerol-3-phosphatase (Gpp). Since Syn2973 lacks an endogenous *gpp* gene, we first introduced *gpp2* from *S. cerevisiae*, generating strain Gly-Strain1 (**Fig. 2B**). Gly-Strain1 produced only 8.63 mg L^-1^ glycerol when cultured from an initial OD_750_ = 0.04 after 96 h (**Fig. S1A**). To enhance precursor flux, we expressed several heterologous *gpd* genes, including *Scegpd1* (*S. cerevisiae*), Fu-SceGpd1-Sce-Gpp2 (a fusion enzyme *Scegpd1 with Scegpp2*), the bifunctional *Cregpd2* (*Chlamydomonas reinhardtii*), *EcogpsA* (*E. coli*), *SyugpsA* (Syn2973), and *BsugpsA* (*Bacillus subtilis*), generating strains Gly-Strain2 to Gly-Strain7 (**Fig. 2B**). Among them, Gly-Strain2 yielded the highest glycerol titer of 134.46 mg L^-1^, followed by Gly-Strain7, respectively (**Fig. S1A**). Given that *Scegpd1* is NADH-dependent whose content is ∼10 fold lower than NADPH in cyanobacteria^21^, *BsugpsA* was further introduced into Gly-Strain2 to form Gly-Strain8, which produced 167.61 mg L^-1^ glycerol after 96 h (**Fig. S1A**). Since the growth state of the strain in different conditions can influence glycerol production (**Fig. S1B**), cells in the stationary phase (OD_750_ = 1, **Fig. S1C**) were used for all subsequent glycerol quantification. Further, we found that genomic integration sites and replicate plasmid expression had negligible effects on growth and glycerol yield (**Fig.S1D** and **S1E**). Alternatively, we found light intensity played a major role. Under medium light (ML, 300 µmol photons·m^-2^·s^-1^), Gly-Strain8 produced 592.16 mg L^-1^ glycerol. Low light (LL, 100 µmol photons·m^-2^·s^-1^) reduced productivity, while high light (HL, 500 µmol photons·m^-2^·s^-1^) imposed stress and decreased yield (**Fig. 2C**).

**Fig. 2.**
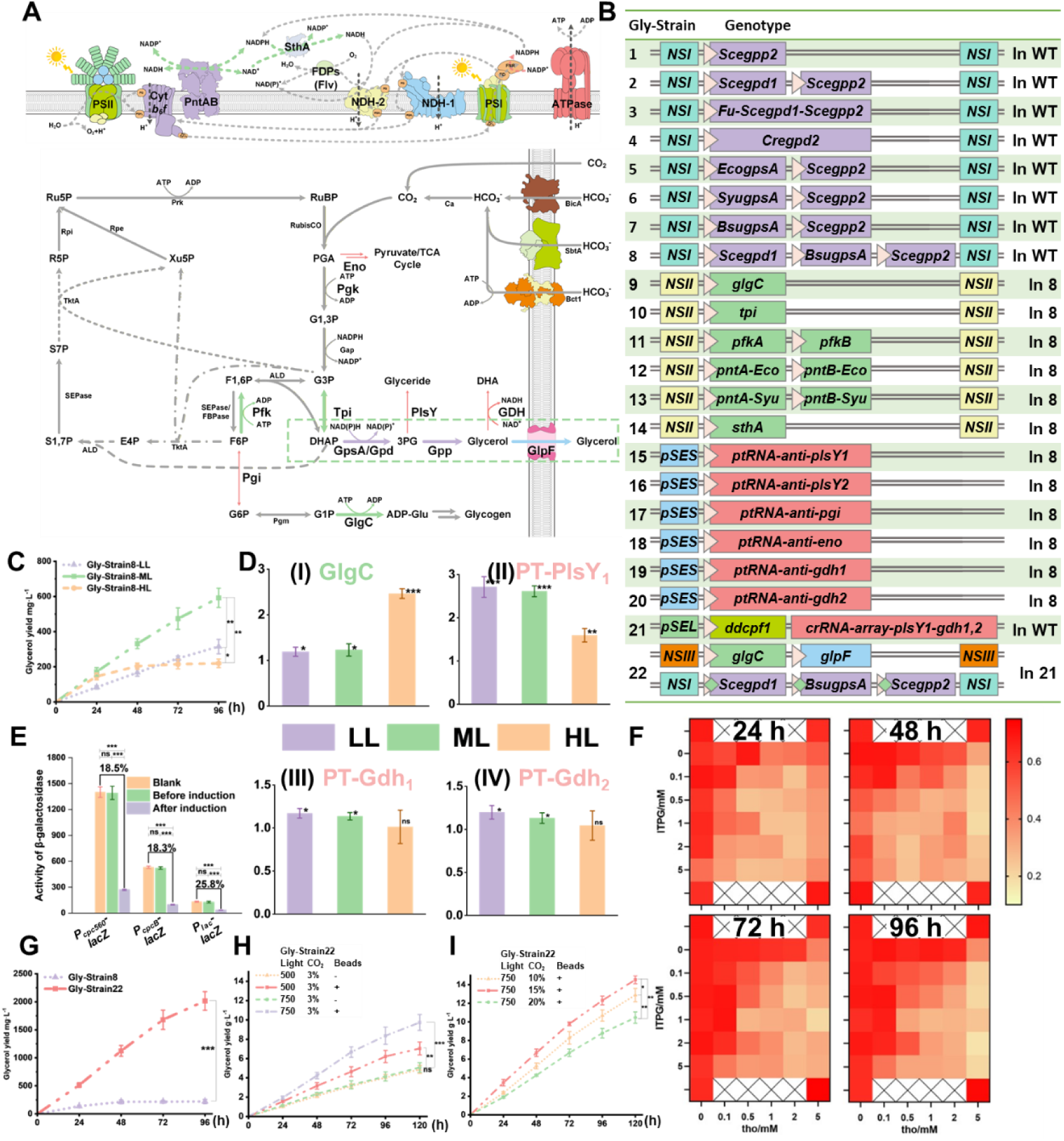
Optimization of the glycerol production in Syn2973. The error bar represents the standard deviation of the three biological replicates for each sample. *P < 0.05, **P < 0.01, ***P < 0.001, statistical analysis was performed using unpaired Student’s t-test. **A)** Schematic of the pathways related to glycerol synthesis and secretion in Syn2973. **B)** Genotype of the strains constructed in this section. **C)** Glycerol production in Gly-Strain8 under different light intensities (LL: 100 µmol photons·m^-2^·s^-1^, ML: 300 µmol photons·m^-2^·s^-1^, HL: 500 µmol photons·m^-2^·s^-1^) **D)** Relative glycerol production in Gly-Strain9 **(i)**, Gly-Strain15 **(ii)**, Gly-Strain19 **(iii)**, and Gly-Strain20 **(iv)** to that in Gly-Strain8 under different light intensities (LL: 100 µmol photons·m^-2^·s^-1^, ML: 300 µmol photons·m^-2^·s^-1^, HL: 500 µmol photons·m^-2^·s^-1^). **E)** Evaluation of the effect of CRISPRi-based gene silence before and after induction. LacZ was used as a reporter, whose expression was controlled under strong, middle and weak promoters (P_cpc560_, P_cpcB_ and P_lac_, respectively). **F)** Optimization of the concentration of inducers (IPTG and theophylline) for CRISPRi-based gene silence. LacZ was used as a reporter, whose expression was controlled under the strong promoter P_cpc560_. **G)** Glycerol production in Gly-Strain8 and Gly-Strain22 in shaking flasks under HL conditions (500 µmol photons·m^-2^·s^-1^). **H)** Glycerol production in Gly-Strain8 and Gly-Strain22 in photoreactors under 3% CO_2_ and light illumination (500 and 750 µmol photons·m^-2^·s^-1^). Beads represented 5 mm glass beads were added to reduce optical path length. **I)** Glycerol production in Gly-Strain8 and Gly-Strain22 in photoreactors under 10%, 15% and 20% CO_2_ and 750 µmol photons·m^-2^·s^-1^ light intensities.

To further boost glycerol synthesis, genes associated with carbon flux or redox balance were overexpressed or repressed in Gly-Strain8. Overexpressed genes included *glgC*, *tpi*, *pfkAB*, *pntAB*, and *sthA* (**Fig. 2A**), generating strains Gly-Strain9 to Gly-Strain14. Repressed genes by PT-RNA^22^ (**Fig. S1F**) included *plsY1*, *plsY2*, *pgi*, *eno*, *gdh1*, and *gdh2*, yielding Gly-Strain15 to Gly-Strain20 (**Fig. 2B**). Compared with Gly-Strain8, *glgC* overexpression (Gly-Strain9) enhanced light tolerance and improved glycerol yield by 2.46-fold under HL conditions (**Fig. 2D (i)**, **Fig. S1G**). Repression of *plsY1* (Gly-Strain14) increased glycerol yield by 2.71-fold (under LL) and 2.61-fold (under ML) (**Fig. 2D (ii)**), while inhibition of *gdh1* and *gdh2* (responsible for glycerol oxidation to DHA) slightly increased glycerol accumulation (**Fig. 2D (iii)** and **(iv)**). Because PlsY1 participates in lipid biosynthesis related to membrane metabolism, its repression impaired early growth (**Fig. S1H**). To address the issue and enable multiplex gene regulation, we employed a ddCpf1-based CRISPRi system with optimized inducible control. Six IPTG-inducible promoters combined with a theophylline-responsive riboswitch were tested to minimize ddCpf1 leakiness and a feedback crRNA (induced by IPTG) targeting *lacI* was added to enhance inducibility (**Fig. S2A, S2B** and **Fig. 2E**). Using *lacZ* as a reporter, combined induction with 2 mM IPTG and 2 mM theophylline achieved strong and stable repression (**Fig. 2F**). Introducing this system into WT (Gly-Strain21) enabled simultaneous repression of *plsY1*, *gdh1*, and *gdh2* without growth defects in uninduced cultures (**Fig. S1H**). Based on Gly-Strain21, expressions of *Scegpd1*, *Scegpp2*, and *BsugpsA* were also placed under the theophylline riboswitch, while *glgC* was overexpressed, and *glpF* was introduced to enhance glycerol export. The resulting strain, Gly-Strain22 (**Fig. 2B**), exhibited markedly increased glycerol titers upon induction, reaching up to 2.02 g L^-1^ under HL (**Fig. 2G**, **S2C**). Then, we optimized photobioreactor conditions to maximize production. Firstly, optimizing the standard BG11 medium with CD medium improved the growth of Gly-Strain22 under 3% CO_2_ aeration and 500 µmol photons·m^-2^·s^-1^ light intensities (**Fig. S2D**). Although cell density increased >10-fold with 3% CO_2_ aeration compared with shake-flask culture, glycerol yield did not scale proportionally (**Fig. 2H**). Based on the phenotypic differences of the engineered strain in aerated and shake flask cultures (**Fig. S2E**), we hypothesized that light attenuation became the primary bottleneck. To address the issue, 5 mm glass beads were added to reduce optical path length (**Fig. S2E**) and growth of Gly-Strain22 under various CO_2_ concentrations (1%-15%) and light intensities (500-750 µmol photons·m^-2^·s^-1^) was evaluated (**Fig. S2F-S2I**). As shown in **Fig. 2H**, **2I** and **S2J**, under 750 µmol photons·m^-2^·s^-1^ light intensity and 15% CO_2_, the highest titer reached 14.53·g L^-1^ after 120 h, representing the highest reported glycerol production in an engineered cyanobacterium to date.

### Construction of glycerol conversion modules for efficient production of 1,3-propanediol and 3-hydroxypropionic acid

As shown in **Fig. 3A**, glycerol can be dehydrated to 3-hydroxypropionaldehyde (3-HPA) by glycerol dehydratase and subsequently converted to either 1,3-propanediol (1,3-PDO) via 3-HPA reductase or to 3-hydroxypropionate (3-HP) via 3-HPA dehydrogenase. Because 3-HPA exhibited strong cytotoxicity to Syn2973 at concentrations above 150 mg·L^-1^ (**Fig. S3A**), we first screened highly active 3-HPA reductases and dehydrogenases. The tested reductases included *E. coli* YqhD and *Klebsiella pneumoniae* DhaT, yielding strains HPA-Strain1 and HPA-Strain2 (**Fig. 3B**). The tested dehydrogenases included Aox (*Pseudomonas putida*), AloD (*Pseudomonas sp.*), KgsadH (*Azospirillum brasilense*) and its mutant KgsadH^E120Q/P219A^ (KgsadHm), AldH (*E. coli*), and GabD4 (*Cupriavidus necator*) and its mutant GabD4^E209Q/E269Q^(GabD4m), generating HPA-Strain3 to HPA-Strain9 (**Fig. 3B**). Among them, HPA-Strain1 (containing YqhD) reduced 106.9 mg·L^-1^ 3-HPA within 24 h, while HPA-Strain9 (containing GabD4^E209Q/E269Q^) oxidized 104.0 mg·L^-1^ under the same conditions (**Fig. 3C**).

**Fig. 3.**
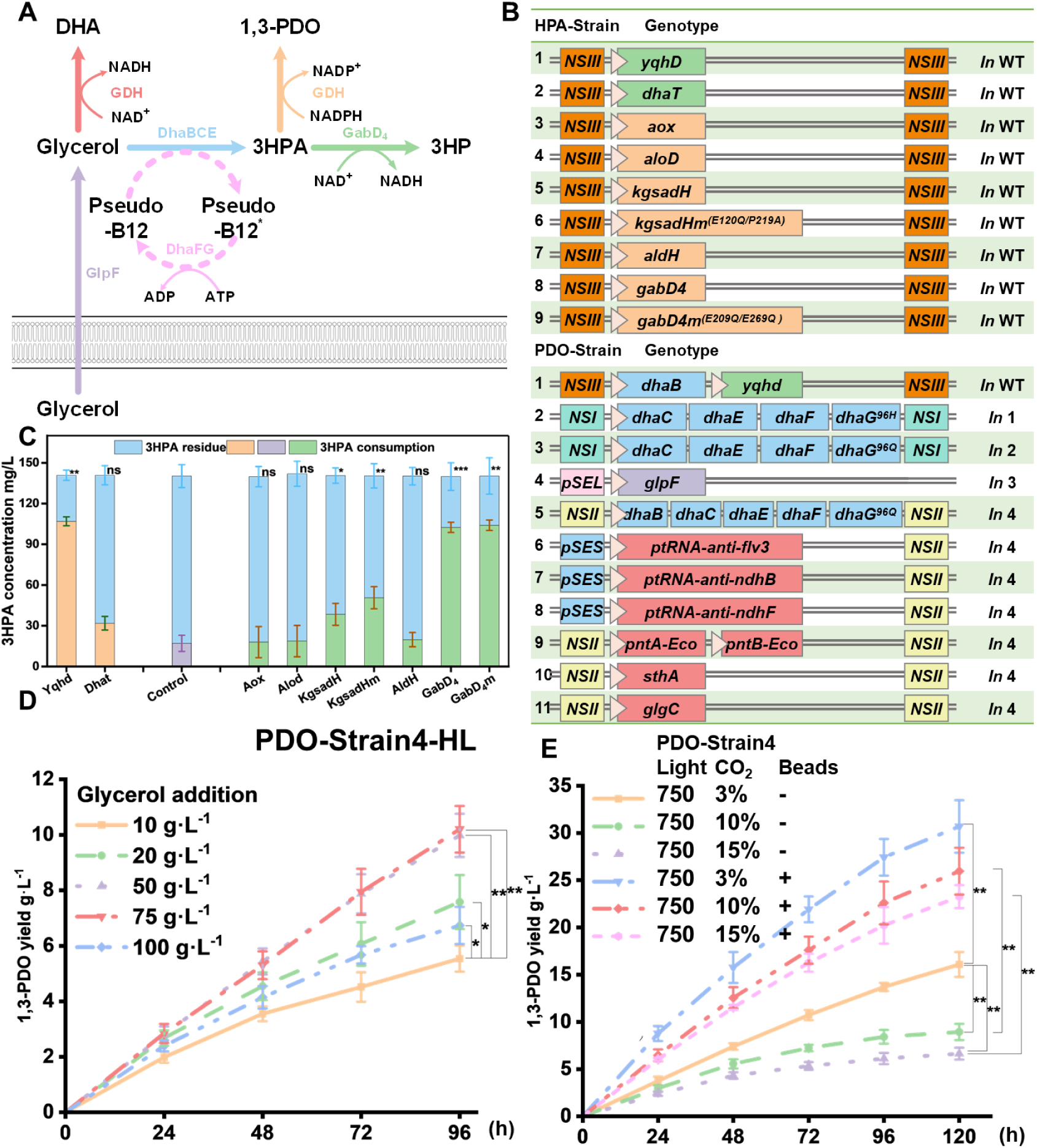
Optimization of the 1,3-PDO production using glycerol in Syn2973. The error bar represents the standard deviation of the three biological replicates for each sample. *P < 0.05, **P < 0.01, ***P < 0.001, statistical analysis was performed using unpaired Student’s t-test. **A)** Schematic of synthesis pathway of 1,3-PDO 3-HP, and DHA based on glycerol. **B)** Genotype of the strains constructed in the section related to 3HPA consumption and 1,3-PDO production. **C)** Results of the screened 3-HPA reductase and 3-HPA dehydrogenase. The blue column represents the residual amount of 3-HPA, while the purple, orange and green columns respectively indicate the consumption of 3-HPA under the action of blank control, 3-HPA reductases and dehydrogenases. **D)** 1,3-PDO production in PDO-Strain4 in shaking flasks under HL (500 µmol photons·m^-2^·s^-1^) supplemented with different concentrations of glycerol (10-100 g·L^-1^). **E)** 1,3-PDO production in PDO-Strain4 in photoreactors under 750 µmol photons·m^-2^·s^-1^ supplemented with different concentrations of glycerol and CO_2_ (3%-15%).

Building on YqhD, we introduced *K. pneumoniae* glycerol dehydratase complex DhaBCE together with its re-activating enzyme DhaFG (variants DhaG^96H^ and DhaG^96Q^), resulting in PDO-Strain2 (DhaBCEFG^96H^) and PDO-Strain3 (DhaBCEFG^96Q^) (**Fig. 3B**). When supplemented with 10 g·L^-1^ glycerol, PDO-Strain3 showed a higher production of 5.54 g·L^-1^ (72.8 mM) within 96 h (**Fig. S3B**). Introducing glycerol facilitator GlpF (PDO-Strain4) further improved production by 10.6% (**Fig. S3B**). Under HL (500 μmol photons·m^-2^·s^-1^) and 50 g·L^-1^ glycerol, PDO-Strain4 reached 9.98 g·L^-1^ 1,3-PDO (**Fig. 3D**, **S3C**, and **S3D**). Notably, in photobioreactors with 3% CO_2_ aeration, 750 µmol photons·m^-2^·s^-1^ light intensities and glass beads for enhanced light transmission, the maximum titer increased to 30.70 g·L^-1^ (**Fig. 3E**, **S3E**). Attempts to further enhance productivity by duplicating the DhaBCEFG^96Q^ module (PDO-Strain5) or supplementing cofactors (vitamin B_12_, Co^2+^) yielded no additional improvement (**Fig. S3F**). We next modified redox and carbon flux pathways to improve reducing force and energy balance, including *flv3*, *ndhB* or *ndhF* repression as well as *glgC, sthA* or *pntAB* overexpression, respectively (PDO-Strain6 to PDO-Strain11, **Fig. 3B**). Only PDO-Strain10, overexpressing *sthA*, showed transient improvement in 1,3-PDO production (**Fig. S3G**), but rapid cell death followed (**Fig. S3H**). Hence, PDO-Strain4 was used for subsequent experiments.

For 3-HP production, DhaB and GabD4m were combined with *K. pneumoniae* DhaCEFG^96Q^, yielding 3HP-Strain2 (**Fig. 4A**). When fed with 10 g·L^-1^ glycerol, 3HP-Strain2 could only produce 3-HP under LL or ML (100 or 300 µmol photons·m^-2^·s^-1^) in shaking flasks, whose viability lost under HL conditions (500 µmol photons·m^-2^·s^-1^) after 24 h (**Fig. S4A**). We attributed this to (i) toxic 3-HPA accumulation (**Fig. S4B**) due to mismatched reaction rates between glycerol dehydration and 3-HPA dehydrogenation and (ii) NAD⁺ depletion during GabD4-mediated oxidation. As proof, induced expression of DhaB (to control the glycerol dehydration) via the theophylline-inducible promoter (3HP-Strain3) prolonged the 3-HP production under HL (**Fig. S4C**). To address the issue, we found introduction of GlpF (3HP-Strain4) alleviated 3-HPA toxicity (**Fig. S4B**), likely due to partial efflux of 3-HPA through GlpF channels, as evidenced by decreased 3-HPA tolerance in WT-GlpF compared with WT. To balance redox metabolism (**Fig. S4D**), 3HP-Strain2 was further engineered to overexpress NADH-consuming or -cycling enzymes, including *flv1,3*, *flv4,2*, *pntAB*, *sthA*, *nox*, *dhaT*, and *yqhD*, or *ndh2* from *Synechocystis* sp. PCC 6803 or *Bacillus subtilis*, resulting in 3HP-Strain5 to 3HP-Strain15 (**Fig. 4A**). Among them, *flv1,3* overexpression (3HP-Strain5) significantly alleviated 3-HPA toxicity (**Fig. S4B**) and improved 3-HP production (**Fig. S4E**). Although YqhD overexpression (3HP-Strain13) reduced toxicity, it also consumed 3-HPA intermediates, leading to lower overall yield. Introduction of GlpF into 3HP-Strain5 created 3HP-Strain16, which was selected for scale-up. With addition of 50 g·L^-1^ glycerol, the strain achieved 6.52 g·L^-1^ 3-HP under shaking flask conditions (500 μmol photons·m^-2^·s^-1^) (**Fig. 4B**) and 9.08 g·L^-1^ using photobioreactors under 750 μmol photons·m^-2^·s^-1^ and 3% CO_2_ (**Fig. 4C**, **S4F**). Further attempts to fine-tune glycerol dehydratase expression (weaker promoters P_lac_ or P_cpcB_; 3HP-Strain17, 3HP-Strain18) or to couple exogenous NADH sinks (Ldh, sAdh, LkAdh; 3HP-Strain19 to 3HP-Strain21) showed no further improvement (**Fig. S4G**, **S4H**). In this case, we speculated that the toxicity of 3-HP might be another major factor (**Fig. S4I**), as our previous study also showed that exogenous addition of only 0.016% (v/v) 3-HP was sufficient to inhibit the growth of *Synechocystis* sp. PCC 6803^23^. Alternatively, we applied a semi-continuous cultivation regime, replacing half or total of the culture daily while feeding glycerol continuously (**Fig. 4D (i)**). Specifically, based on the results of **Fig. 4D** (**ii**), when 5 g·L^-1^ or 7.5 g·L^-1^ glycerol was added daily, half of the culture medium was replaced each day (cells were retained by centrifugation, and 3HP-Strain16 achieved productivities of 4.16 g·L^-1^·day^-1^and 6.43 g·L^-1^·day^-1^, with corresponding glycerol conversion rates of 85.1% and 87.7%, respectively (**Fig. 4D** (**iii**), (**iv)**). When 10 g·L^-1^ or higher glycerol was added daily, the entire medium needed to be replaced; under 10 g·L^-1^ addition, the strain produced 6.30 g·L^-1^·day^-1^ 3-HP, corresponding to a conversion rate of 64.4% (**Fig. 4D** (**v**), (**vi**)). All the semi-continuous cultivation regime could last for at least 5 d.

**Fig. 4.**
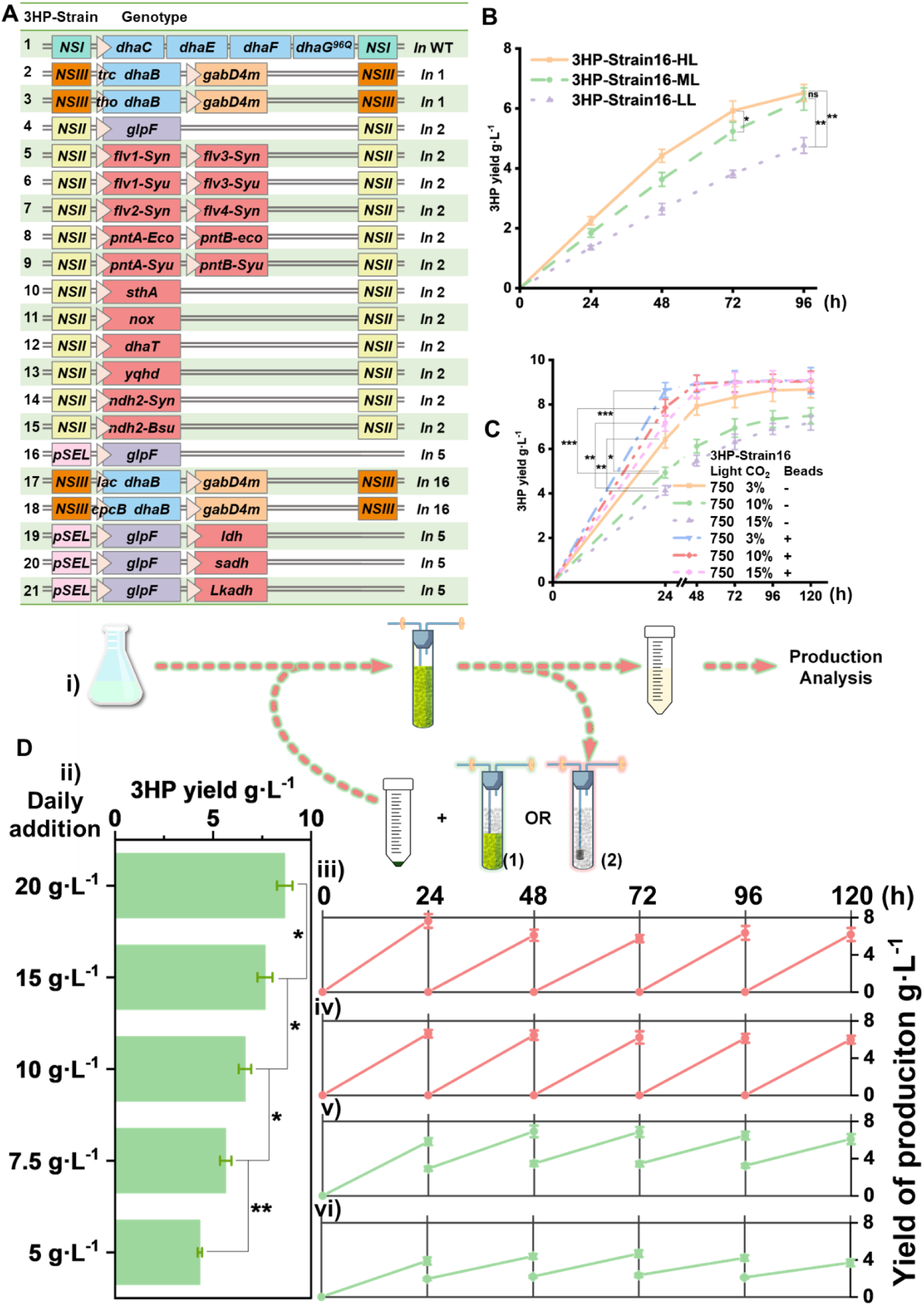
Optimization of the 3-HP production using glycerol in Syn2973. The error bar represents the standard deviation of the three biological replicates for each sample. *P < 0.05, **P < 0.01, ***P < 0.001, statistical analysis was performed using unpaired Student’s t-test. **A)** Genotype of the strains constructed in the section related to 3HP production. **B)** 3-HP production in 3HP-Strain16 cultured in shaking flasks under LL, ML, and HL (100, 300 and 500 µmol photons·m^-2^·s^-1^).**C)** 3-HP production in 3HP-Strain16 cultured in photoreactors under HL (750 µmol photons·m^-2^·s^-1^) supplemented with 3%, 10%, and 15% CO_2_. **D)** Semi-continuous cultivation for 3HP-Strain16 enabled efficient 3-HP production. (i) Schematic of the semi-continuous cultivation regime. (ii) Daily 3-HP production when supplemented with different concentrations of glycerol in photoreactors (750 µmol photons·m^-2^·s^-1^ supplemented with 3% CO_2_). the 3-HP productivity of the semi-continuous cultivation regime when replacing half of the culture daily while feeding glycerol (5 g·L^-1^ (iii) or 7.5 g·L^-1^ (iv)) continuously. the 3-HP productivity of the semi-continuous cultivation regime when replacing total of the culture daily while feeding glycerol (10 g·L^-1^ (v) or 15 g·L^-1^ (vi)) continuously.

### Construction of glycerol conversion modules for efficient production of DHA

Glycerol can be oxidized to DHA via one-step glycerol dehydrogenase-mediated reaction (**Fig. 3A**). We evaluated a series of dehydrogenases, including the endogenous *SyuGldA*, *EcoGldA* (*E. coli*), *KpnGldA* and *KpnDhaD* (*K. pneumoniae*), *SceGcy1* (*S. cerevisiae*), and *GoxSldAB* (*G. oxydans*), generating DHA-Strain1 to DHA-Strain6 (**Fig. 5A**). Among these, DHA-Strain2 (*EcoGldA*) and DHA-Strain3 (*KpnGldA*) showed the highest catalytic efficiency (**Fig. S5A** and **S5B**). Co-expression of both *EcoGldA* and *KpnGldA* yielded DHA-Strain7, and further introduction of the glycerol facilitator *GlpF* produced DHA-Strain8 (**Fig. 5A**). However, DHA-Strain8 did not exhibit proportional increases in DHA production with glycerol consumption. In shake-flask cultures, DHA accumulation remained low (212.59 mg·L^-1^) despite much glycerol utilization, and in photoreactors, 14.86 mM glycerol was converted into only 4.82 mM DHA (**Fig. 5B**). To improve redox balance and photosynthetic efficiency, we introduced the same genetic modifications tested in the 3-HP pathway—targeting *flv1,3*, *flv4,2*, *nox*, *ndh2*, *pntAB*, and *sthA* genes from *Synechocystis* sp. PCC 6803, Syn2973, *E. coli*, or *B. subtilis*—to generate DHA-Strain9 to DHA-Strain17. None of these strains showed enhanced DHA accumulation (**Fig. S5C**). Given the chemical reactivity of DHA^24^, we speculated that DHA might be unstable or further metabolized in Syn2973. Indeed, stability tests revealed that DHA rapidly degraded under alkaline conditions but remained relatively stable at neutral or slightly acidic pH (**Fig. 5C**). Because cyanobacterial cytosol is typically alkaline (pH > 8) to support photosynthetic reactions^25^, DHA degradation may occur intracellularly. Buffering the culture medium to pH 7.5 partially stabilized DHA in the supernatant (**Fig. 5C**). Moreover, the presence of an endogenous *gdh* gene suggested that an uncharacterized DHA catabolic pathway might exist. Together, these findings indicate that Syn2973 may not be suitable as a chassis for DHA production.

**Fig. 5.**
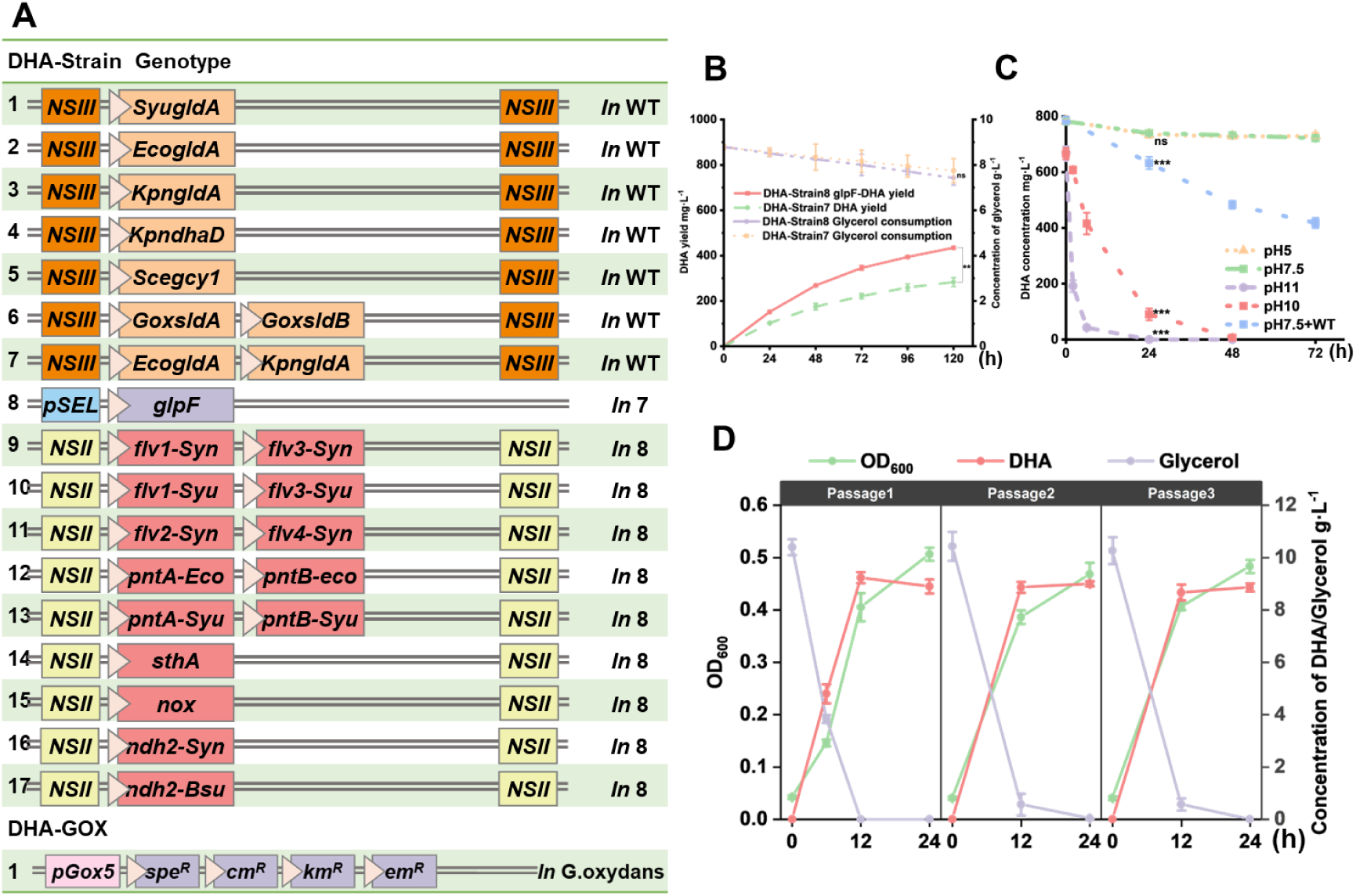
Optimization of the 3-HP production using glycerol in Syn2973. The error bar represents the standard deviation of the three biological replicates for each sample. *P < 0.05, **P < 0.01, ***P < 0.001, statistical analysis was performed using unpaired Student’s t-test. **A)** Genotype of the strains constructed in the section related to DHA production. **B)** DHA production and glycerol consumption in DHA-Strain1 cultured in in photoreactors under 750 µmol photons·m^-2^·s^-1^ supplemented with 3% CO_2_. **C)** Stability evaluation of DHA under medium with different pH and Syn2973 culture. **D)** Growth, glycerol consumption, and DHA production of DHA-GOX1 cultured for three successive generations in CD^M2^ medium preconditioned by Syn2973.

As an alternative, we employed *G. oxydans* 621H, a strain that naturally oxidizes glycerol to DHA with minimal nutrient requirement^26^. The main challenge was to establish a compatible co-culture medium with the glycerol-producing Syn2973. We compared the compositions of CD medium for Syn2973 with GMM and MSM media for *G. oxydans* 621H, identifying key differences in nitrogen sources and cofactors such as nicotinic acid, p-aminobenzoic acid, pantothenic acid, and glutamate (**Fig. S5D**). Based on CD medium, we formulated a modified medium (CD^M1^) by supplementing these components, buffering with TES to pH 7.0, and adjusting NH_4_Cl concentration to 0.25 g·L^-1^. We next introduced an empty plasmid conferring shared antibiotic resistance into *G. oxydans*, generating DHA-GOX1 (**Fig. 5A**). DHA-GOX1 grew well in CD^M1^ and efficiently converted glycerol to DHA (**Fig. S5E**). To further simplify the medium, we removed nicotinic acid, p-aminobenzoic acid, pantothenic acid, and glutamate from CD^M1^, generating CD^M2^. However, we found that the DHA-GOX1 strain could sustain normal growth and production in CD^M2^ for only one generation (**Fig. S5F**). Assuming that Syn2973 <u>or its cell debris and intracellular contents</u> could supply the added cofactors <u>or their derivatives</u> in co-culture, we removed nicotinic acid, p-aminobenzoic acid, pantothenic acid, and glutamate to create a simplified CD^M2^ medium. On one hand, the modified medium supported normal growth of Syn2973 (**Fig. S5G**). On the other hand, when DHA-GOX1 was cultivated in CD^M2^ preconditioned by Syn2973, it maintained robust growth and DHA synthesis (**Fig. 5D**), reaching a maximal glycerol conversion rate up to 87.9%. Notably, the growth and production efficiency of DHA-GOX1 cultivated in CD^M2^ was comparable to that cultivated in GMM medium (**Fig. S5H**).

### Linking Glycerol Production with Downstream Conversion Expands CO2 Bioconversion to 1,3-PDO, 3-HP, and DHA

In this study, we evaluated two strategies for coupling glycerol production and conversion modules. The first was a sequential process, in which the glycerol-producing strain was cultured and removed after glycerol accumulation, followed by inoculation with the conversion strain (**Fig. 6A (ii)**). The second was a co-culture strategy, where the production and conversion modules were mixed at defined ratios and cultivated in a single system (**Fig. 6A (iii)**). As a reference, we also constructed complete CO_2_-to-product pathways in a single Syn2973 strain for 1,3-PDO, 3-HP, and DHA (**Fig. 6A (i)**). Using the most efficient pathway combinations identified in this study, we generated strains PDO-Strain12, 3HP-Strain22, and DHA-Strain18 (**Fig. 6B**). Accordingly, the production of 1,3-PDO, 3-HP, and DHA reached 3.24 g·L^-1^, 5.42 g·L^-1^, and 441.38 mg·L^-1^ using photoreactors (15% CO_2_ and illumination of 750 μmol photons·m^−2^·s^−1^), respectively (**Fig. 6C**, **6D**, and **6E**).

**Fig. 6.**
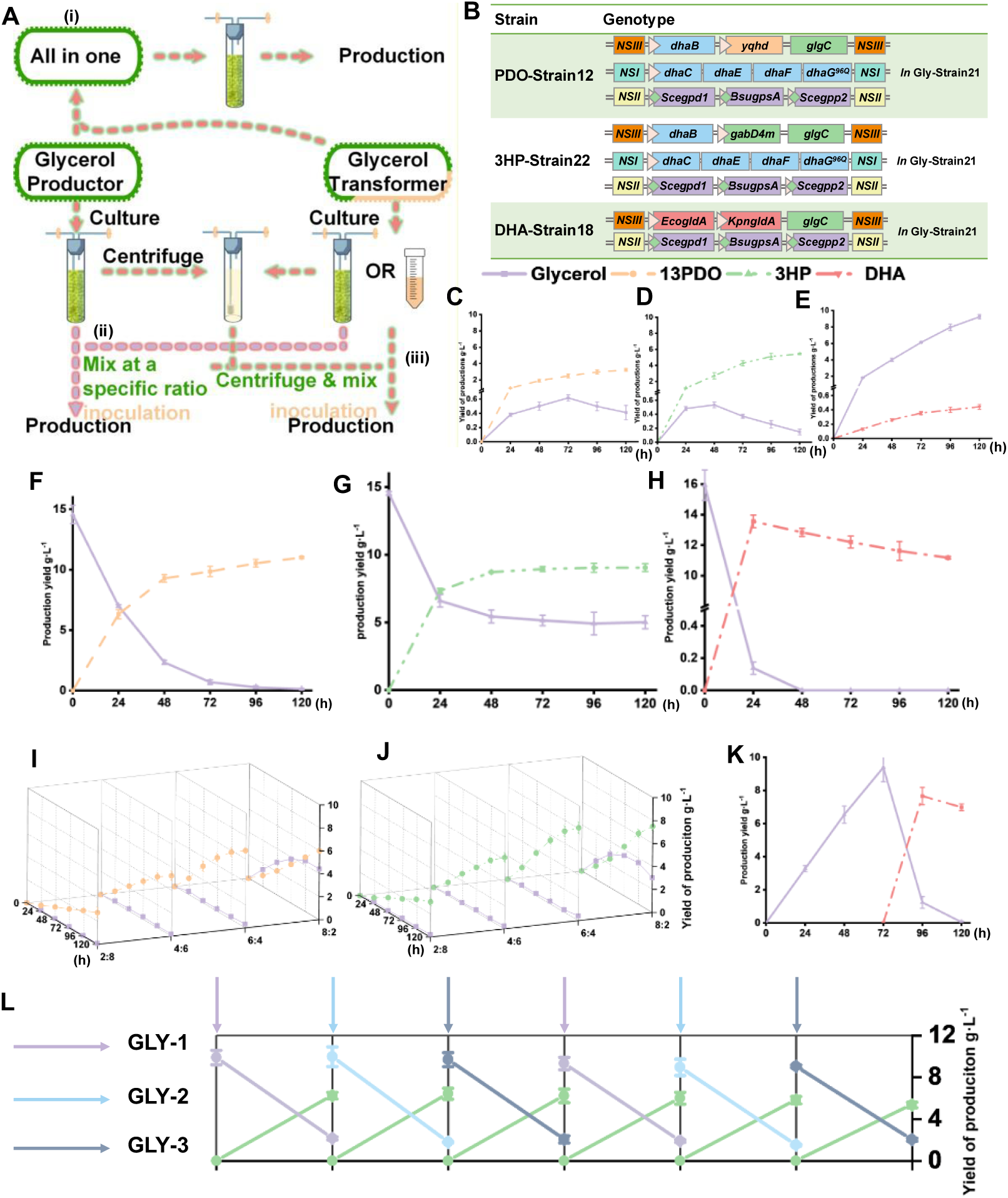
Optimization of the co-culture systems composed of the glycerol biosynthesis and conversion module. The error bar represents the standard deviation of the three biological replicates for each sample. *P < 0.05, **P < 0.01, ***P < 0.001, statistical analysis was performed using unpaired Student’s t-test. **A)** (i) schematic illustration of the production of 1,3-PDO, 3-HP and DHA directly from CO_2_. (ii) Schematic illustration of the first co-culture strategy, in which the glycerol-producing module was removed after glycerol accumulation and subsequently replaced with the conversion module to transform glycerol into 1,3-PDO, 3-HP, or DHA. (iii) Schematic illustration of the second co-culture strategy, where the glycerol-producing and conversion modules were co-inoculated at defined ratios to enable simultaneous glycerol production and conversion. **B)** Genotypes of strains constructed for direct biosynthesis of 1,3-PDO, 3-HP, or DHA from CO_2_. **C–E)** Glycerol accumulation and production of 1,3-PDO (**C**), 3-HP (**D**), and DHA (**E**) in PDO-Strain12, 3HP-Strain22, and DHA-Strain18, respectively, cultured in photobioreactors (3% CO_2_, 750 µmol photons·m^-2^·s^-1^). **F–H)** Glycerol accumulation and production of 1,3-PDO (**F**), 3-HP (**G**), and DHA (**H**) using the co-culture strategy; for **H**, the conversion module used was *G. oxydans* strain DHA-GOX1. **I-K)** Glycerol accumulation and production of 1,3-PDO (**I**), 3-HP (**J**), and DHA (**K**) using the co-culture strategy; for **K**, the conversion module used was DHA-GOX1, which was introduced 72 h after initiation of the glycerol-producing module. **L)** Semi-continuous conversion of glycerol to 1,3-PDO using Gly-Strain22 and PDO-Strain4. Specifically, PDO-Strain4 was inoculated into the culture supernatant of Gly-Strain22 grown for 72 h; after 24 h of conversion, PDO-Strain4 cells were collected and transferred into fresh 72-h Gly-Strain22 cultures. This process was stably repeated for at least six cycles.

For the sequential modular system, the glycerol-producing strain Gly-Strain22 was first cultivated and induced for 120 h, accumulating approximately 14.73 g·L^-1^ (160 mM) glycerol. The cells were then removed, and the culture medium was inoculated with PDO-Strain4, 3HP-Strain16, DHA-Strain8 or DHA-GOX1. Within 120 h, glycerol was successfully converted into 11.00 g·L^-1^ 1,3-PDO, 9.05 g·L^-1^ 3-HP, 435.36 mg·L^-1^ DHA (using DHA-Strain8), and 13.52 g·L^-1^ DHA (using DHA-GOX1), respectively (**Fig. 6F**, **6G**, **S6A**, and **6H**).

For the co-culture system, Gly-Strain22 was mixed with PDO-Strain4, 3HP-Strain16, DHA-Strain8 or DHA-GOX1 at different inoculation ratios. For both 1,3-PDO and 3-HP production, the optimal inoculation ratio of 6:4 (production: conversion) yielded 6.74 g·L^-1^ 1,3-PDO and 8.01 g·L^-1^ mM 3-HP, respectively (**Fig. 6I** and **6J**). Co- culture of Strain22 and DHA-Strain8 with a ratio of 1:9 reached 393.62 mg·L^-1^ DHA production (**Fig. S6B**). In contrast, co-cultivation of Gly-Strain22 with DHA-GOX1 at an initial OD_600_ of 0.1 led to DHA synthesis but caused bleaching and death of Gly-Strain22 after 48 h (**Fig. S6C**). Toxicity analysis revealed that 5 g·L^-1^ DHA imposed severe growth inhibition on Syn2973 (**Fig. S6D**). To mitigate this effect, DHA-GOX1 was introduced 48 h after glycerol accumulation, which successfully produced 7.66 g·L^-1^ DHA (85.03 mM) (**Fig. 6K**). Based on this optimized configuration, we further tested the system under continuous and semi-continuous cultivation conditions using the production of 1,3-PDO as a proof of concept. First, we determined the maximum daily glycerol conversion capacity of PDO-Strain4. Culture supernatants from Gly-Strain22 grown for one, two, or three days were collected (**Fig. S6E**) and used to inoculate PDO-Strain4. The results showed that glycerol was efficiently converted under all conditions (**Fig. S6F**). Based on these findings, we designed a semi-continuous co-culture strategy for glycerol-to-1,3-PDO conversion. Specifically, PDO-Strain4 was inoculated daily into the culture supernatant of Gly-Strain22 that had been cultivated for three days, enabling at least six days of continuous production. Under these conditions, the system achieved an average productivity of 6.02 g·L^-1^·day^-1^ (79.05 mM·day^-1^), with a glycerol to 1,3-PDO conversion efficiency of 75.1% (**Fig. 6L**). Using the final semi-continuous cultivation system as an example, we estimated the net CO_2_ fixation of the entire production process. Because CO_2_ was the sole carbon source, the evaluation followed the equation: E_CO2_ = [-(W_CB_ × C_C_/12) - (m_product_/M_product_ × C_product_)] × M_CO2_^20^. In this system, the final cell dry weights of the three glycerol-producing strains and the 1,3-PDO–producing strain were 5.25, 5.11, 5.53, and 8.05 g·L^-1^, respectively. After six days of operation, the 1,3-PDO titer reached 6.02 g·L^-1^, with 1.92 g·L^-1^ residual glycerol. Notably, current process of 1,3-PDO production release CO_2_ at 2.5 to 6.7 kg CO_2_ eq/kg 1,3-PDO^27^. In contrast, our system produced 6.02g·L^-1^ 1,3- propanediol while fixing 20.68 kg CO_2_ per liter, corresponding to a net carbon fixation efficiency of 3.43 kg CO_2_ eq/kg 1,3-PDO.

## Discussion

Rewiring photosynthetic cyanobacteria for efficient CO_2_-to-chemical conversion has remained challenging, with productivities far below those of heterotrophic chassis such as *E. coli* or *K. pneumoniae* ^28, 29^. Even with extensive pathway reconstruction, cyanobacterial strains typically generate only trace-level titers of target molecules, reflecting fundamental constraints in redox supply, metabolite toxicity, and cellular resource allocation. Here we introduce a modular metabolic architecture that breaks from the classical single-cell chassis: instead of forcing photosynthetic cells to perform both carbon fixation and complex chemical transformations, we segregate these metabolic functions and use glycerol as a dynamically exchanged metabolite. This division-of-labor strategy bypasses long-standing bottlenecks that have limited the synthesis of highly reduced chemicals such as 1,3-PDO and 3-HP, which previously reached only millimolar levels in cyanobacteria^30, 31^. mainly constrained by limited reducing power and intermediate toxicity. By decoupling CO_2_ fixation and product formation, our glycerol-mediated platform effectively bypasses these constraints. The engineered Syn2973 strain achieved over 14 g·L^-1^ glycerol, which is one order of magnitude higher than the best-performing photosynthetic chassis reported so far^32^. Our results demonstrate that glycerol can serve as a robust and redox-efficient carbon relay in photosynthetic metabolism.

Our co-culture configurations highlight the broader potential of both autotroph–autotroph and autotroph-heterotroph partnerships as distributed biomanufacturing systems^20^. Although recent cyanobacteria-bacteria consortia have demonstrated CO_2_-to-lactate, 2,3-butanediol, or 3-HP production ^18, 33^, these systems are typically constrained by competition for light, carbon flux, or metabolic space. By contrast, using glycerol as a nonvolatile, rapidly diffusible intermediate enables clean metabolic segregation: photosynthetic cells dedicate resources to carbon assimilation, while partner strains execute high-flux reductive or oxidative transformations. This design minimized metabolic interference and supported potential full conversion of glycerol into 1,3-PDO, 3-HP, and DHA.

Because modules and partners can be interchanged, the platform can be readily adapted for other chemistries, positioning metabolite-mediated communication as a generalizable strategy for photosynthetic biomanufacturing. Conceptually, this work establishes a blueprint for modular CO_2_ utilization in which pathway orthogonalization and stable intermediate shuttling jointly overcome the intrinsic limitations of single-cell systems. With further integration of spatial and temporal control, such architectures could achieve higher productivities while maintaining strict carbon neutrality. Beyond 1,3-PDO and 3-HP, this framework offers a foundation for producing a wide portfolio of C_3_-C_4_ chemicals, moving toward sunlight-powered, carbon-negative chemical factories.

## Methods

### Strains and culture conditions

Wild-type *Synechococcus elongatus* UTEX 2973 (Syn2973) and its engineered derivatives were cultured in standard BG11 medium (pH 7.5) under shaking conditions (200 rpm) at 37 °C with continuous illumination (∼500 μmol photons·m^-2^·s^-1^) using an orbital shaker incubator (HNYC-202T, Ounuo, Tianjin, China) ^34^. For solid cultivation, cells were streaked onto BG11 agar plates and incubated at 37 °C under 300 μmol photons m^-2^·s^-1^ (SPX-250B-G, Boxun, Shanghai, China). *Escherichia coli* strains were cultivated in LB medium at 37 °C, while *Gluconobacter oxydans* 621H was grown at 37 °C in either sorbitol-based medium (73 g·L^-1^ sorbitol, 18.4 g·L^-1^ yeast extract, 1.5 g·L^-1^ (NH_4_)_2_SO_4_, 1.5 g·L^-1^ KH_2_PO_4_, 0.47 g·L^-1^ MgSO_4_·7H_2_O) or defined salt medium. Antibiotics were added as needed: for Syn2973 and *G. oxydans*, 20 μg·mL^-1^ kanamycin, 20 μg·mL^-1^ chloramphenicol, 50 μg·mL^-1^ spectinomycin and/or 50 μg·mL^-1^ erythromycin were used; for *E. coli*, 100 μg·mL^-1^ ampicillin, 50 μg·mL^-1^ kanamycin, 50 μg·mL^-1^ chloramphenicol, 100 μg·mL^-1^ spectinomycin and/or 200 μg·mL^-1^ erythromycin. For induction, IPTG or theophylline was added at concentrations specified in the Results section. Cell densities were measured as OD_750_ for Syn2973 and OD_600_ for *E. coli* and *G. oxydans* using a microplate reader (TECAN Infinite E Plex).

### Plasmid and strain construction

*E. coli* DH5α was used for plasmid propagation. Primers and synthetic DNA fragments were synthesized by GENEWIZ (Suzhou, China). Plasmids were constructed using seamless cloning (ClonExpress MultiS One Step Cloning Kit, Vazyme, Nanjing, China) or Golden Gate assembly. Details of primers, templates, and assembly strategies are listed in **Table S2**; sequences of synthetic fragments are provided in **Supplementary Table S3**. Syn2973 was transformed via triparental conjugation, while *G. oxydans* was transformed via electroporation. Strains used in this study are listed in Supplementary **Table S1**.

### β-galactosidase assay

β-Galactosidase activity was determined as previously described. Syn2973 cells in exponential phase were collected at a culture volume equivalent to OD_750_×1. Cells were centrifuged and resuspended in 1 mL Z buffer (60 mM Na_2_HPO_4_, 40 mM NaH₂PO₄, 10 mM KCl, 1 mM MgSO_4_, 40 mM β-mercaptoethanol), followed by the addition of 50 μL 0.1% SDS and 50 μL chloroform to lyse the cells. The reaction was initiated by adding 200 μL ONPG (4 g·L^-1^) as substrate and incubated at 30 °C, 750 rpm for 3 min. The reaction was terminated with 500 μL 1 M Na_2_CO_3_, and OD_420_ was measured to calculate Miller Units using the formula: Miller = 1000 × OD_420_ / 3^13^.

### Photobioreactor cultivation

Syn2973 was cultured in 100 mL flat-bottom glass tubes (30 mm diameter) with a working volume of 60 mL. BG11 medium was replaced with 5×BG or CD medium when specified. The culture was bubbled with a gas mixture of air and CO_2_ (flow rate = culture volume, i.e., 1 vvm) controlled by a mass flow controller. Initial OD_750_ was set to 0.1. Light intensity was programmed as follows: 0-12 h, 100 μmol photons·m^-2^·s^-1^; 12-24 h, 200 μmol photons·m^-2^·s^-1^; 24-36 h, <u>300</u> photons·m^-2^·s^-1^; and after 36 h, 500 μmol photons·photons·m^-2^·s^-1^. If higher light intensities were required, adjustments were made after 48 h. Temperature was maintained using a water circulation system.

### Quantitative real-time PCR (qRT-PCR)

Cells were harvested after 48 h of cultivation at OD_750_×volume = 1 (equals 1.45 x 10^8^ cells). Total RNA was extracted using the Direct-zol^TM^ RNA MiniPrep Kit (Zymo Research, CA, USA) following the manufacturer’s instructions. RNA concentration and quality were assessed via Nanodrop 2000 and agarose gel electrophoresis. First-strand cDNA was synthesized using the RevertAid First Strand cDNA Synthesis Kit (Thermo Fisher Scientific) with 500 ng RNA as template. qPCR was performed using PowerUp SYBR Green Master Mix (Thermo Fisher Scientific) in a 10 μL reaction system containing 5 μL master mix, 3 μL ddH_2_O, 1 μL cDNA (1:1000 dilution), and 0.5 μL each of forward and reverse primers (see **Table S2**). The reactions were run on a StepOnePlus™ Real-Time PCR System (Applied Biosystems). Relative gene expression was calculated using the 2⁻^ΔΔCT^ method ^35^ with 16S rRNA as internal control. Each sample was measured in triplicate.

### Metabolite quantification via HPLC

Glycerol, 1,3-propanediol (1,3-PDO), 3-hydroxypropionic acid (3-HP), 3-hydroxypropionaldehyde (3-HPA), and dihydroxyacetone (DHA) concentrations in culture supernatants were measured by HPLC (Agilent 1260) equipped with an Aminex HPX-87H column (300 mm × 7.8 mm, Bio-Rad) and operated at 65 °C with 5 mM H_2_SO_4_ as mobile phase at a flow rate of 0.6 mL·min⁻¹. UV-Vis and refractive index (RID) detectors were used simultaneously. Injection volume was 10 μL. All reported concentrations are means of three biological replicates. Glycerol and 1,3-PDO were detected exclusively by RID. 3-HP and DHA, which overlap with glycerol in RID, were quantified by UV signal and subtracted from total RID to derive glycerol concentration. 3-HPA, having only a weak UV signal and poor peak shape, was quantified by reacting equal volumes of sample and Fehling’s reagent, incubating at 60 °C for 5 min, then measuring the <u>increase</u> in 3-HP concentration via HPLC to indirectly calculate 3-HPA content. Representative chromatograms are provided in Supplementary **Fig. S7**.

## Conclusions

In summary, this study establishes a modular photosynthetic platform that enables efficient CO_2_-to-chemical conversion through glycerol-mediated metabolic division. By integrating optimized glycerol biosynthesis in Syn2973 with downstream conversion modules for 1,3-PDO, 3-HP, and DHA, we demonstrate that functional coupling between autotrophic and heterotrophic partners can overcome the long-standing limitations of redox imbalance, product toxicity, and pathway competition in cyanobacterial biosynthesis. The glycerol-mediated system not only achieves record-level production in photosynthetic cyanobacteria but also introduces a generalizable design principle for distributed carbon metabolism. This work thus provides a conceptual and technological foundation for scalable, carbon-negative biomanufacturing, highlighting the potential of modular consortia as a next-generation strategy for synthetic biology. The framework presented here can be extended to a wide array of reductive and oxidative chemical transformations, paving the way toward sunlight-driven, sustainable chemical synthesis.

## Data availability statement

All the data have been included in the supplementary file.

## SUPPLEMENTARY DATA

Supplementary Data is available online.

