## Supplemental Figures and Tables for "Synthetic Modular Cyanobacterial Consortium Enables Carbon-Negative Production of Glycerol and Derivatives"

*[1] School of Synthetic Biology and Biomanufacturing, Tianjin University, Tianjin 300072, P.R. China；[2] Frontier Science Center for Synthetic Biology and Key Laboratory of Systems Bioengineering, Ministry of Education of China, Tianjin 300072, P.R. China; [3] Center for Biosafety Research and Strategy, Tianjin University, Tianjin 300072, P.R. China; [4] State Key Laboratory of Synthetic Biology, Tianjin University, Tianjin, 300072, P.R. China; [5] Haihe Laboratory of Sustainable Chemical Transformations, Tianjin 300192, China.*

† These two authors contributed equally to the paper.

* To whom all correspondence should be addressed:

Prof. Dr. Weiwen Zhang
School of Synthetic Biology and Biomanufacturing
Tianjin University, Tianjin 300072, P.R. China


Prof. Dr. Lei Chen
School of Synthetic Biology and Biomanufacturing
Tianjin University, Tianjin 300072, P.R. China


Prof. Dr. Tao Sun
School of Synthetic Biology and Biomanufacturing
Tianjin University, Tianjin 30072, P.R. China.


**Running title**: Synthetic Modular Cyanobacterial Consortium

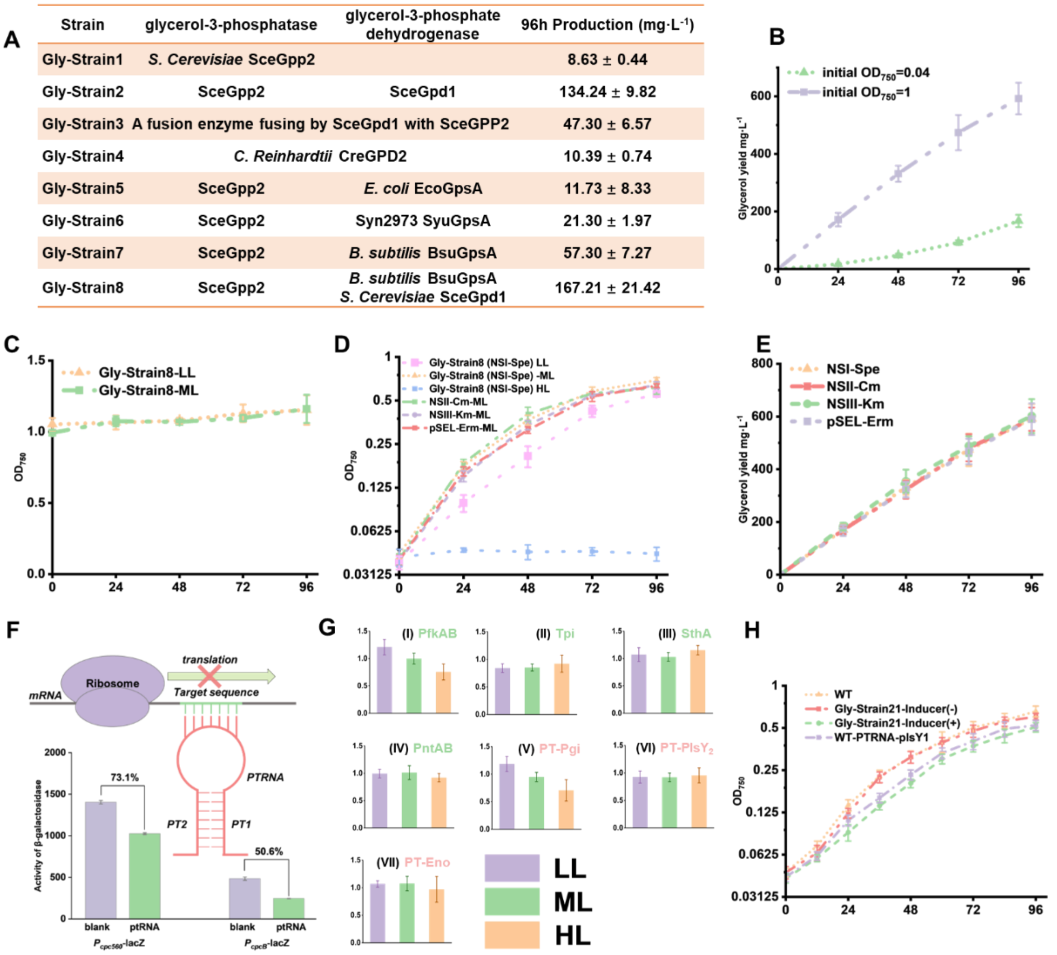

| Strain | glycerol-3-phosphatase | glycerol-3-phosphate dehydrogenase | 96h Production (mg·L-1) |
|---|---|---|---|
| Gly-Strain1 | *S. Cerevisiae* SceGpp2 | | 8.63 ± 0.44 |
| Gly-Strain2 | SceGpp2 | SceGpd1 | 134.24 ± 9.82 |
| Gly-Strain3 | A fusion enzyme fusing by SceGpd1 with SceGPP2 | | 47.30 ± 6.57 |
| Gly-Strain4 | *C. Reinhardtii* CreGPD2 | | 10.39 ± 0.74 |
| Gly-Strain5 | SceGpp2 | *E. coli* EcoGpsA | 11.73 ± 8.33 |
| Gly-Strain6 | SceGpp2 | Syn2973 SyuGpsA | 21.30 ± 1.97 |
| Gly-Strain7 | SceGpp2 | *B. subtilis* BsuGpsA | 57.30 ± 7.27 |
| Gly-Strain8 | SceGpp2 | *B. subtilis* BsuGpsA *S. Cerevisiae* SceGpd1 | 167.21 ± 21.42 |

**Fig. S1. A)** Glycerol production of strains Gly-Strain1 to Gly-Strain8 after 96 h of cultivation. **B)** Glycerol production at different inoculation densities (OD = 0.04 and OD = 1). These results indicate that the physiological state of the cells affects glycerol yield; therefore, an inoculation density of OD = 1 was used for all subsequent assays. **C)** Consistent with panel B, using Gly-Strain8 as an example, biomass remained nearly unchanged under low and medium light intensities when an inoculation density of OD = 1 was used for glycerol production. **D–E)** Growth and glycerol production of Gly-Strain8 derivatives expressing the glycerol biosynthesis module integrated into different genomic loci (NSI, NSII, NSIII), antibiotic selection marker or maintained on a shuttle plasmid (pSEL), showing no significant effect on glycerol yields. **F)** Schematic diagram of the PTRNA interference system and its validation using *lacZ* as a reporter. Expression driven by either the high-strength promoter $P_{cpc560}$ or the medium-strength promoter $P_{cpcB}$ demonstrated effective suppression. **G)** Supporting data for Fig. 1D: Relative glycerol titers in shake-flask cultures under low, medium, and high light after overexpression of *pfkAB*, *tpi*, *sthA*, or *pntAB*, or suppression of *pgi*, *plsY2*, or *eno*, compared with Gly-Strain8. **H)** Growth curves of Gly-Strain8, Gly-Strain14, and Gly-Strain22 under induced and non-induced conditions. The data demonstrates that inducible repression of *plsY1* allows normal growth to be restored in the absence of induction.

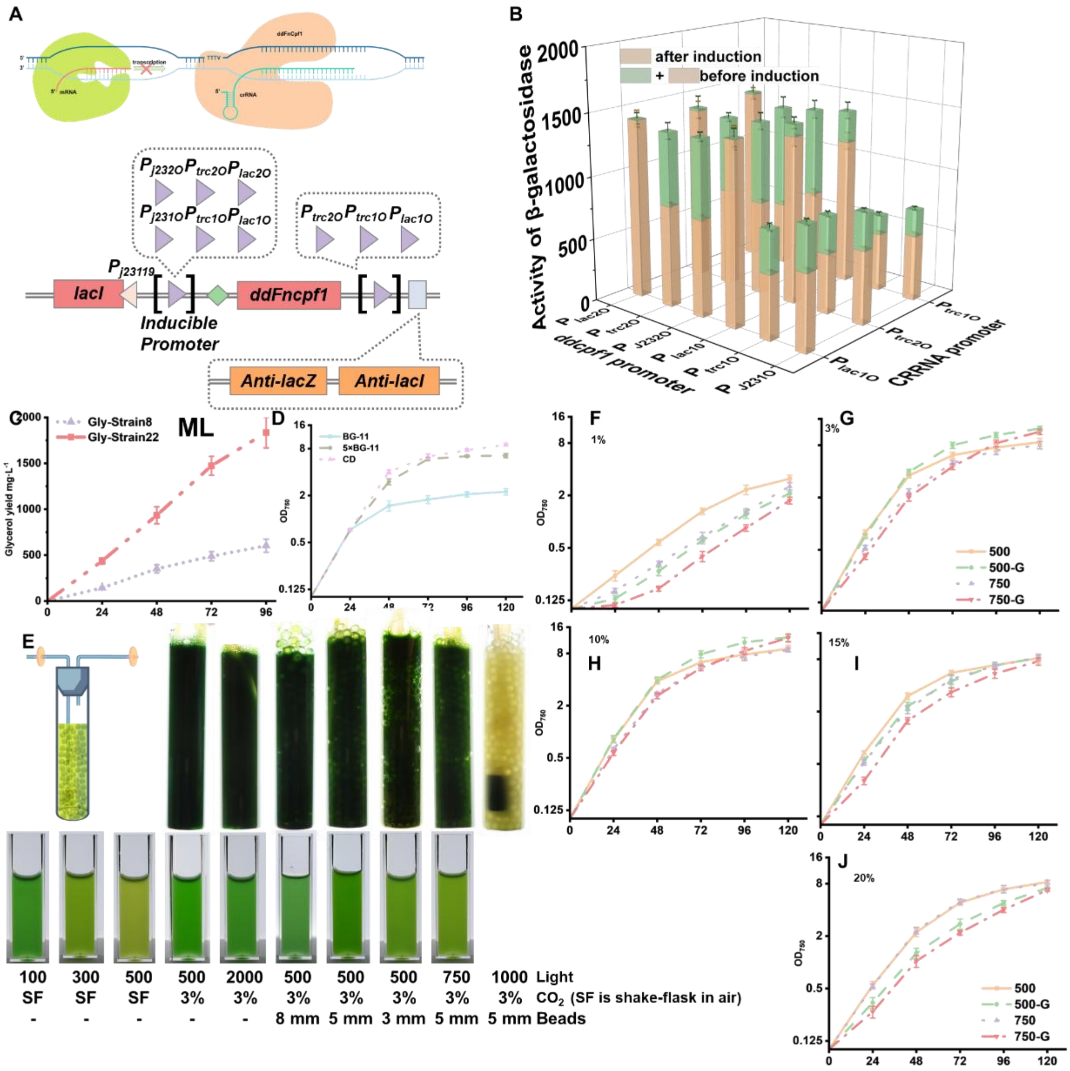

**Fig. S2. A)** Schematic of the ddFnCpf1-based CRISPRi interference system and the optimization strategy implemented in this study. **B)** Initial testing of the system using β-galactosidase (LacZ) as a reporter gene. In the original design, *lacI* was driven by promoter $P_{j23119}$, ddFnCpf1 by $P_{trc2O}$, and the crRNA by $P_{trc1O}$ (2O and 1O indicate the number of LacI operator sites). IPTG was used as the inducer. However, this configuration failed to yield detectable repression of LacZ (Fig. S2B). To enhance repression efficiency, an additional crRNA targeting *lacI* was introduced into the crRNA array. Although this improved repression of the target gene, substantial repression was also observed in the absence of inducer. Subsequently, various promoter combinations for driving ddFnCpf1 and crRNA expression were evaluated, but none produced the desired inducible repression profile. To overcome this limitation, a theophylline-responsive riboswitch was incorporated to further regulate ddFnCpf1 expression. The $P_{trc1O}$–thoE*–$P_{trc1O}$ configuration showed the most effective performance, and this optimized system was validated across targets with varying expression levels (see Fig. 2E in the main text). **C)** Glycerol production of Gly-Strain8 and Gly-Strain23 under ML (300 μmol photons·m$^{-2}$·s$^{-1}$). **D)** Growth curves of Gly-Strain23 in photobioreactors under different medium conditions (3% $CO_2$ and 750 μmol photons·m$^{-2}$·s$^{-1}$). **E)** Phenotypic comparison of cultures grown in shake flasks versus photobioreactors. In shake flasks, strong light penetration results in yellowing under high-light conditions, while in photobioreactors without glass beads, cultures remained green even under 2000 μmol photons·m$^{-2}$·s$^{-1}$, indicating insufficient light penetration. Addition of glass beads restored the expected yellow phenotype, confirming improved light distribution. **F–J)** Growth curves of Gly-Strain23 in photobioreactors under 1% $CO_2$ (F), 3% $CO_2$ (G), 10% $CO_2$ (H), 15% $CO_2$ (I), and 20% $CO_2$ (J) at light intensities of 500 and 750 μmol photons·m$^{-2}$·s$^{-1}$.

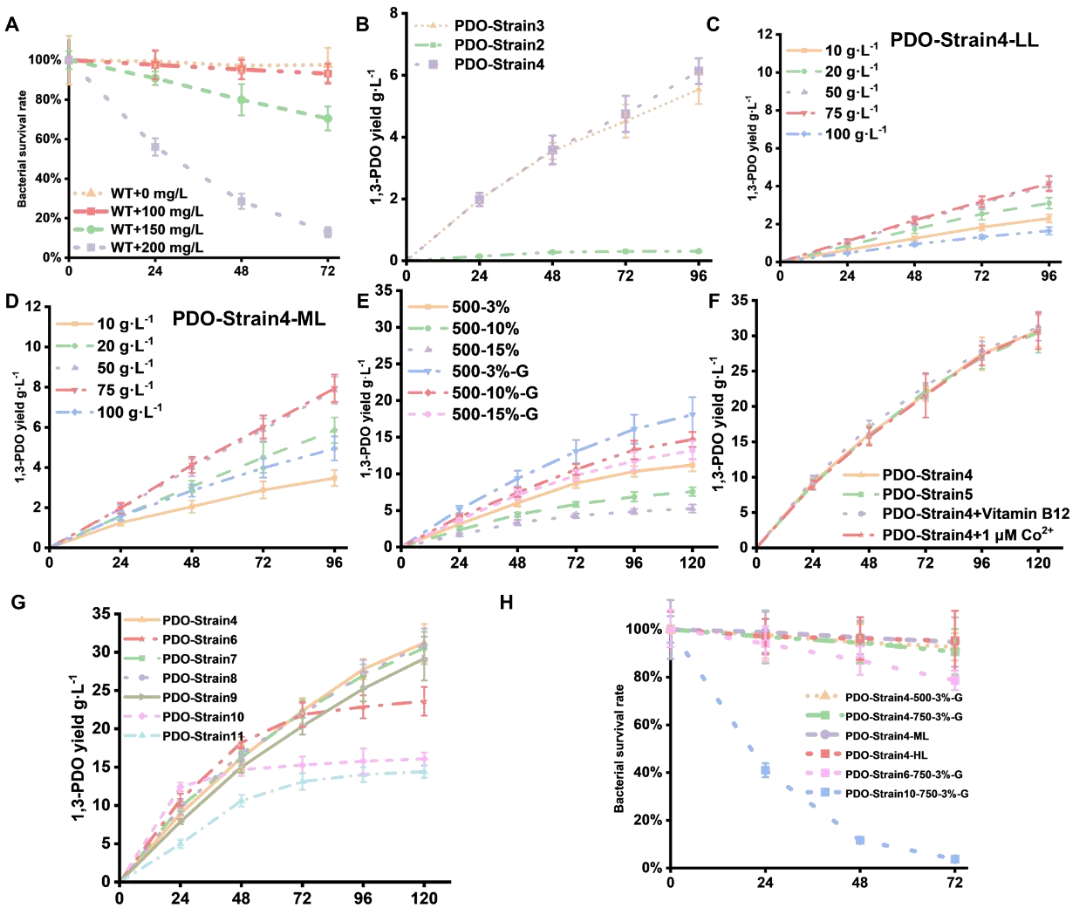

**Fig. S3. A)** Cell viability of the wild-type (WT) strain under different concentrations of exogenously added 3-HPA. **B)** 1,3-PDO production in PDO-Strain2, PDO-Strain3 and PDO-Strain4 cultured in shake flasks under high-light conditions (500 μmol photons·$m^{-2}\cdot s^{-1}$). **C)** and **D)** 1,3-PDO production in PDO-Strain4 cultured in shake flasks under low-light (100 μmol photons·$m^{-2}\cdot s^{-1}$) and medium-light (300 μmol photons·$m^{-2}\cdot s^{-1}$) conditions, respectively. **E)** 1,3-PDO production in PDO-Strain4 grown in photobioreactors at 500 μmol photons·$m^{-2}\cdot s^{-1}$ under different $CO_2$ concentrations. **F)** 1,3-PDO production in PDO-Strain5 and PDO-Strain4 under standard conditions or supplemented with 3 μM vitamin $B_{12}$ or 1 μM $Co^{2+}$. Cultivation was performed in photobioreactors at 3% $CO_2$ and 750 μmol photons·$m^{-2}\cdot s^{-1}$. **G)** 1,3-PDO production in PDO-Strain6, …, PDO-Strain11 under photobioreactor conditions (3% $CO_2$ and 750 μmol photons·$m^{-2}\cdot s^{-1}$). **H)** Cell viability of PDO-Strain6, …, PDO-Strain11 under the same cultivation conditions.

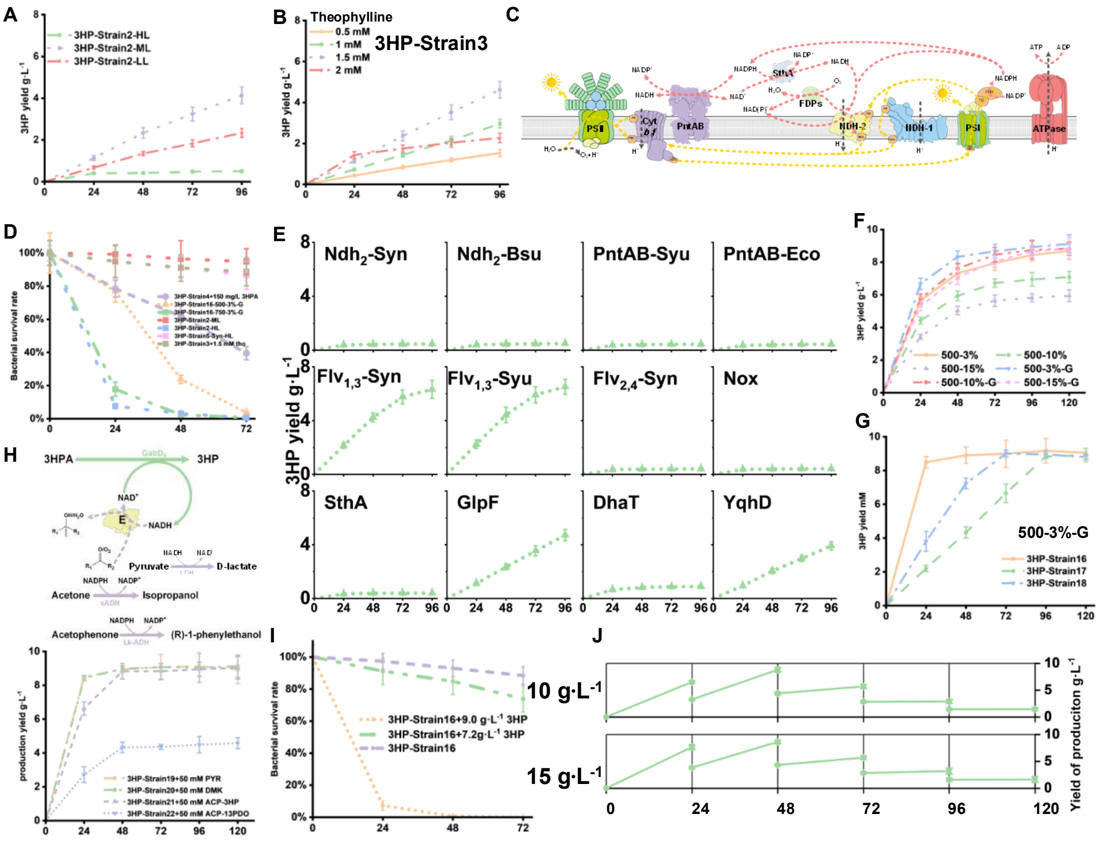

**Fig. S4. A)** Time-course profiles of 3-HP production by 3HP-Strain2 under different light intensities (LL: 100 μmol photons·m$^{-2}$·s$^{-1}$; ML: 300 μmol photons·m$^{-2}$·s$^{-1}$; HL: 500 μmol photons·m$^{-2}$·s$^{-1}$). **B)** 3-HP production of 3HP-Strain3 in response to varying theophylline induction levels. **C)** Schematic overview of energy metabolism–related pathways and genes in the photosynthetic system. **D)** Cell viability of 3HP-Strain2…, 3HP-Strain5 and 3HP-Strain16. **E)** 3-HP production of 3HP-Strain4 to 3HP-Strain15 in shaking flasks under 500 μmol photons·m$^{-2}$·s$^{-1}$. **F)** 3-HP production of 3HP-Strain16 under different $CO_2$ concentrations (3%, 10%, and 15%) at 500 μmol photons·m$^{-2}$·s$^{-1}$ in photobioreactors. **G)** Comparison of 3-HP production in 3HP-Strain16, 3HP-Strain17, and 3HP-Strain18 under photobioreactor conditions (3% $CO_2$ and 750 μmol photons·m$^{-2}$·s$^{-1}$). Strain17 and Strain18 were engineered with promoter substitutions for *dhaB*. **H)** Schematic of additional NADH-balancing pathways introduced through alternative product formation and 3-HP production levels of the corresponding engineered strains (3% $CO_2$ and 750 μmol photons·m$^{-2}$·s$^{-1}$). **I)** Cell viability of 3HP-Strain16 at different exogenous 3-HP concentrations. **J)** Semi-continuous cultivation performance of 3HP-Strain16 with daily addition of 10 g/L or 15 g/L glycerol. Results indicate that with glycerol feeding exceeding 10 g/L and daily replacement of half of the medium, the strain gradually lost its ability to synthesize 3-HP from glycerol starting on day 3.

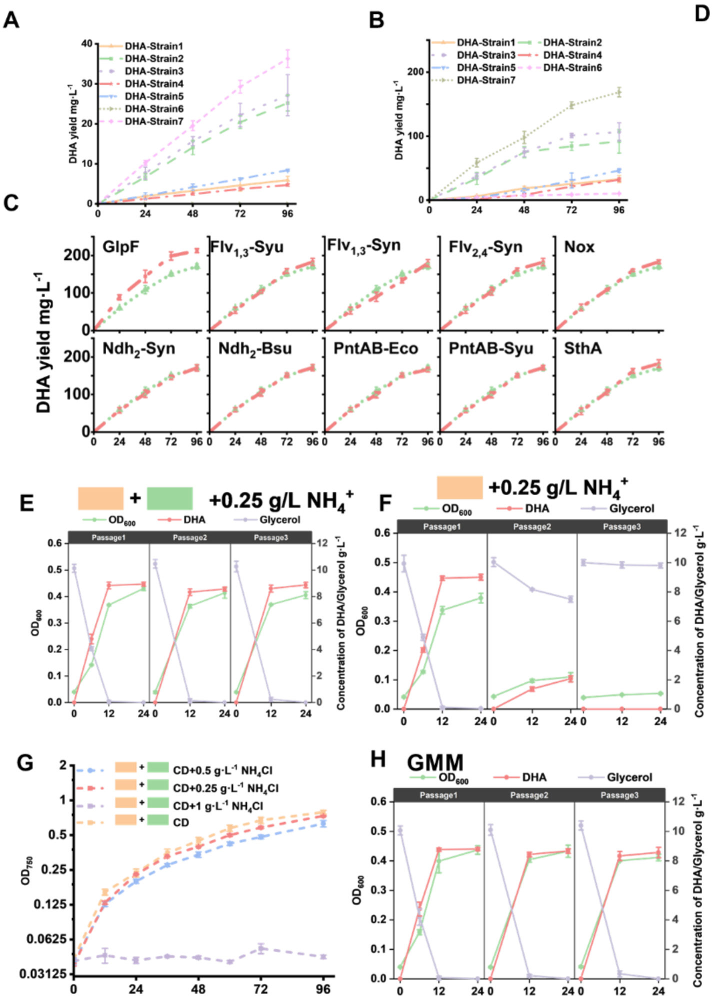

D

| | CD | GMM | Mineral Salt Medium |
|---|---|---|---|
| $KNO_3$ | 50 | | |
| $NaNO_3$ | 50 | | |
| $(NH4)_2SO_4$ | | 7.56 | 15.135 |
| $K_2HPO4$ | | | |
| $KH_2PO4$ | 4 | 7.35 | 16.166 |
| $Na_2HPO4$ | | | 1.409 |
| $MgSO_4$ | 5 | 10.14 | 0.811 |
| $CaCl_2$ | 3 | 0.0476 | 0.061 |
| Citric acid | | | |
| Ferric Ammonium Citrate | | | |
| $FeCl_2$ | | 0.0394 | 0.045 |
| $FeCl_3$ | 0.45 | 0.0185 | |
| EDTA | 0.45 | | 0.103 |
| Nitrilotriacetic acid | | | 0.026 |
| $Na_2CO_3$ | | | |
| $H_3BO_3$ | 0.23 | | 0.016 |
| $MnCl_2$ | 0.045 | 0.0036 | 0.01 |
| $ZnSO_4$ | 0.00385 | | 0.031 |
| $Na_2MoO_4$ | 0.008 | | 0.0003 |
| $(NH_4)_6Mo_7O_{24}$ | | 0.0024 | |
| $CuSO_4$ | 0.0015 | 0.016 | 0.0024 |
| $CoCl_2$ | 0.00085 | 0.0178 | 0.0025 |
| KI | | | 0.0012 |
| Nicotinic acid | | 0.008 | 0.0032 |
| P-aminobenzoic acid | | 0.0729 | 0.0029 |
| Pantothenic acid | | 0.0046 | 0.002 |
| Glutamate | | 10.195 | 1.369 |
| Glycine | | 2.331 | |
| Isoleucine | | 1.601 | |

**Fig. S5. A)** DHA production profiles of DHA-Strain1 to DHA-Strain7 in shake-flask cultivation under 500 μmol photons·m$^{-2}$·s$^{-1}$. **B)** DHA production profiles of DHA-Strain1 to DHA-Strain7 in photobioreactors supplied with 3% $CO_2$ and 750 μmol photons·m$^{-2}$·s$^{-1}$. **C)** DHA production profiles of DHA-Strain8 to DHA-Strain17 in photobioreactors supplied with 3% $CO_2$ and 750 μmol photons·m$^{-2}$·s$^{-1}$. **D)** Comparative analysis of Syn2973 CD medium, *Gluconobacter* GMM medium, and Mineral Salt Medium, followed by formulation of a modified medium capable of supporting co-cultivation of Syn2973 and *Gluconobacter*. Orange-highlighted components represent shared ingredients included at higher concentrations; purple-highlighted components denote ammonium salts, whose concentration was optimized due to ammonium toxicity to Syn2973; green-highlighted components represent unique ingredients from *Gluconobacter* media absent in CD. **E)** Growth, DHA production, and glycerol consumption of *Gluconobacter* cultivated for three consecutive generations in mixed medium CD$^{M1}$. Inclusion of green-highlighted components (panel E) restored stable growth and sustained DHA production. **F)** Growth, DHA production, and glycerol consumption of *Gluconobacter* cultivated for three consecutive generations in mixed medium CD$^{M2}$. In the absence of green-highlighted components (panel E), the strain failed to maintain stable growth and production. **G)** Growth curves of Gly-

Strain22 in $CD^{M2}$ medium supplemented with different concentrations of ammonium sulfate, used to determine optimal ammonium levels for the mixed medium. **H)** Growth, DHA production, and glycerol consumption of *Gluconobacter* cultivated for three consecutive generations in GMM medium.

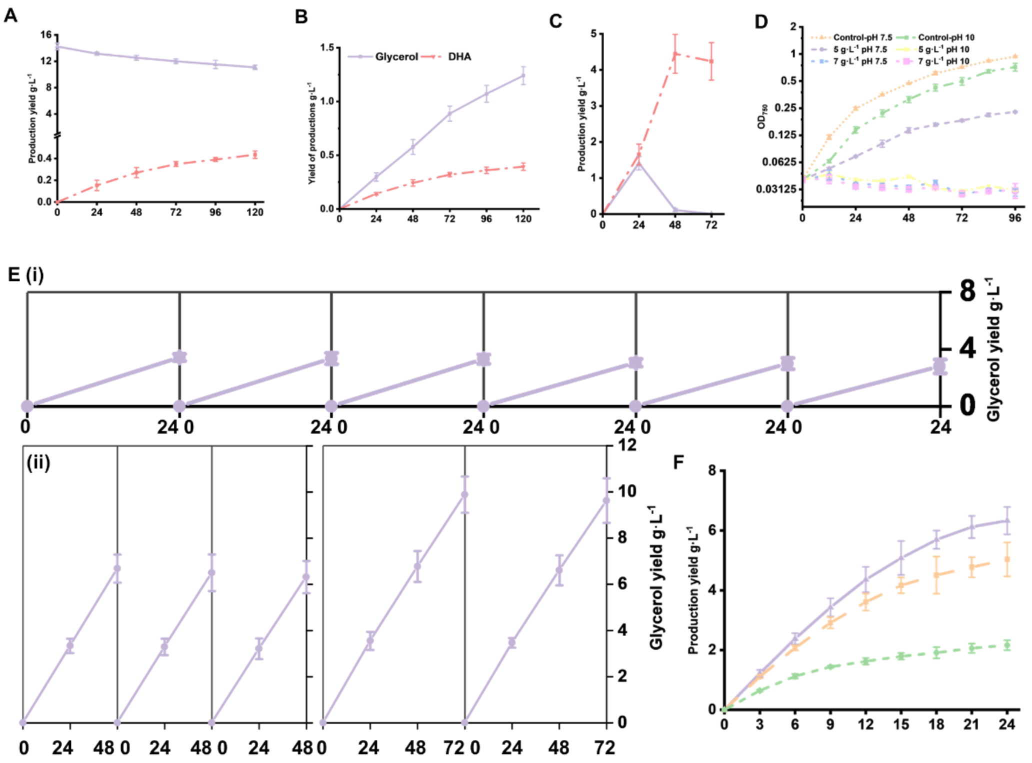

**Fig. S6. A)** Glycerol consumption and DHA production in the co-culture of DHA-Strain18 and Gly-Strain22, corresponding to the mixed-culture mode shown in Fig. 6A (ii) of the main text. **B)** Glycerol consumption and DHA production in the co-culture of DHA-Strain18 and Gly-Strain22, corresponding to the mixed-culture mode shown in Fig. 6A (iii) of the main text. **C)** Glycerol consumption and DHA production in the co-culture of DHA-GOX1 and Gly-Strain22 under the mixed-culture conditions shown in Fig. 6A (iii) of the main text. **D)** Growth curves of Gly-Strain22 under different pH values and varying concentrations of exogenously added DHA. **E)** Glycerol production of Gly-Strain22 when harvested every 24 h (i), 48 h (ii), or 72 h (iii). The 24 h harvesting strategy supports continuous cycling for six rounds, the 48 h strategy for three rounds, and the 72 h strategy for two rounds. **F)** Corresponding to panel **E**: culture supernatants collected after 24 h, 48 h, or 72 h of Gly-Strain22 cultivation were inoculated with PDO-Strain4, and 1,3-PDO production was measured every 3 h over a 24 h period.

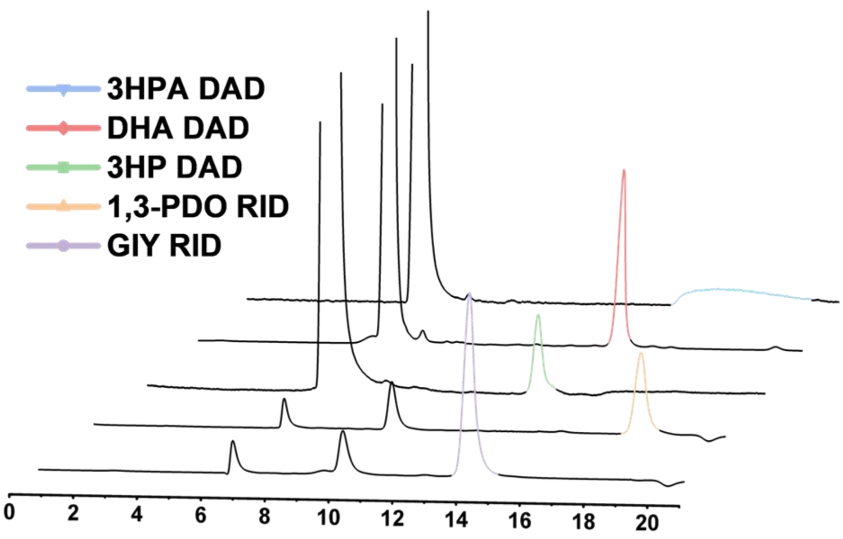

**Fig. S7. HPLC profiles of standards for glycerol, 3-HPA, 3-HP, 1,3-PDO, and DHA dissolved in spent medium from Syn2973 cultures.** Due to the poor peak shape of 3-HPA in this matrix, quantification of 3-HPA in this study was performed by first oxidizing it to 3-HP and subsequently measuring 3-HP.

**Table S1. Strains constructed or used in this study.**

| Strains | Genotype |
|---|---|
| ***E. coli* strains** | |
| DH5a | $F^-$φ80d *lacZ*ΔM15 Δ(*lacZYA-argF*) U169 *endA*1 *recA*1 *hsdR*17 ($r_k^-$,$m_k^+$) *supE*44 , $λ^-$ *thi*-1 *gyrA*96 *relA*1 *pho*A |
| HB101 | *supE*44, Δ(*mcrC-mrr*), *recA*13, *ara*-14, *proA*2, *lacY*1, *galK*2, *rpsL*20, *xyl*-5, *mtl*-1, *leuB*6, *thi*-1 |
| ***g. oxydans* strains** | |
| GOXDHA | pGOX5::pGOX5-spe-cm-km-erm; $spe^R$, $km^R$, $cm^R$ and $em^R$ in wild type Synechococcus elongatus UTEX 2973 |
| **Cyanobacterial strains** | |
| WT | wild type *Synechococcus elongatus* UTEX 2973 |
| WT-LacZ | NSI::pSI-cpc560-lacZ; $spe^R$ in WT |
| Gly-Stran1 | NSI::pSI-cpc560-scegpp2; $spe^R$ in WT |
| Gly-Stran2 | NSI::pSI-cpc560-scegpd1-scegpp2; $spe^R$ in WT |
| Gly-Stran3 | NSI::pSI-cpc560-Fu-scegpd1-scegpp2; $spe^R$ inWT |
| Gly-Stran4 | NSI::pSI-cpc560-cregpd2; $spe^R$ in WT |
| Gly-Stran5 | NSI::pSI-cpc560-ecogpsA-scegpp2; $spe^R$ in WT |
| Gly-Stran6 | NSI::pSI-cpc560-syugpsA-scegpp2; $spe^R$ in WT |
| Gly-Stran7 | NSI::pSI-cpc560-bsugpsA-scegpp2; $spe^R$ in WT |
| Gly-Stran8 | NSI::pSI-cpc560-scegpd1-bsugpsA-scegpp2; $spe^R$ in WT |
| Gly-Stran8-NSII | NSII::pSII-cpc560-scegpd1-bsugpsA-scegpp2; $cm^R$ in WT |
| Gly-Stran8-NSIII | NSIII::pSIII-cpc560-scegpd1-bsugpsA-scegpp2; $km^R$ in WT |
| Gly-Stran8-panL | pSEL-cpc560-scegpd1-bsugpsA-scegpp2; $em^R$ in WT |
| WR-cpcB-LacZ | NSI::pSI-cpcB1-lacZ; $spe^R$ in WT |
| LacZ-PT | NSII::pSII-ptRNA-lacZ; $cm^R$ in WT-lacZ |
| cpcB-LacZ-PT | NSII::pSII-ptRNA-lacZ; $cm^R$ in WT-cpcB-lacZ |
| Gly-Stran9 | NSII::pSII-cpcB-glgC; $cm^R$ in Gly-Stran8 |
| Gly-Stran10 | NSII::pSII-cpcB-tpi; $cm^R$ in Gly-Stran8 |
| Gly-Stran11 | NSII::pSII-cpcB-pfkAB; $cm^R$ in Gly-Stran8 |
| Gly-Stran12 | NSII::pSII-cpcB-pntAB-eco; $cm^R$ in Gly-Stran8 |
| Gly-Stran13 | NSII::pSII-cpcB-pntAB-syu; $cm^R$ in Gly-Stran8 |
| Gly-Stran14 | NSII::pSII-cpcB-sthA; $cm^R$ in Gly-Stran8 |
| Gly-Stran15 | pSES-ptRNA-plsY1; $cm^R$ in Gly-Stran8 |
| Gly-Stran16 | pSES-ptRNA-plsY2; $cm^R$ in Gly-Stran8 |

| | |
|---|---|
| Gly-Stran17 | pSES-ptRNA-pgi; *cm*$^R$ in Gly-Stran8 |
| Gly-Stran18 | pSES-ptRNA-eno; *cm*$^R$ in Gly-Stran8 |
| Gly-Stran19 | pSES-ptRNA-GDH1; *cm*$^R$ in Gly-Stran8 |
| Gly-Stran20 | pSES-ptRNA-GDH2; *cm*$^R$ in Gly-Stran8 |
| LacZ-ddCpf1-Trc2O-Trc1O | pSEL-ddCPF1-anti-lacZ Trc2O-Trc1O; *km*$^R$ *i*n WT-lacZ |
| LacZ-ddCpf1-lacI-Trc2O-Trc1O | pSEL-ddCPF1-anti-lacI-lacZ Trc2O-Trc1O; *km*$^R$ in WT-lacZ |
| LacZ-ddCpf1-lacI-Trc2O-Lac1O | pSEL-ddCPF1-anti-lacI-lacZ Trc2O-Lac1O; *km*$^R$ in WT-lacZ |
| LacZ-ddCpf1-lacI-Trc2O-Trc2O | pSEL-ddCPF1-anti-lacI-lacZ Trc2O-Trc2O; *km*$^R$ in WT-lacZ |
| LacZ-ddCpf1-lacI-Trc1O-Trc1O | pSEL-ddCPF1-anti-lacI-lacZ Trc1O-Trc1O; *km*$^R$ in WT-lacZ |
| LacZ-ddCpf1-lacI-Trc1O-Lac1O | pSEL-ddCPF1-anti-lacI-lacZ Trc1O-Lac1O; *km*$^R$ in WT-lacZ |
| LacZ-ddCpf1-lacI-Trc1O-Trc2O | pSEL-ddCPF1-anti-lacI-lacZ Trc1O-Trc2O; *km*$^R$ in WT-lacZ |
| LacZ-ddCpf1-lacI-Lac2O-Trc1O | pSEL-ddCPF1-anti-lacI-lacZ Lac2O-Trc1O; *km*$^R$ in WT-lacZ |
| LacZ-ddCpf1-lacI-Lac2O-Lac1O | pSEL-ddCPF1-anti-lacI-lacZ Lac2O-Lac1O; *km*$^R$ in WT-lacZ |
| LacZ-ddCpf1-lacI-Lac2O-Trc2O | pSEL-ddCPF1-anti-lacI-lacZ Lac2O-Trc2O; *km*$^R$ in WT-lacZ |
| LacZ-ddCpf1-lacI-Lac1O-Trc1O | pSEL-ddCPF1-anti-lacI-lacZ Lac1O-Trc1O; *km*$^R$ in WT-lacZ |
| LacZ-ddCpf1-lacI-Lac1O-Lac1O | pSEL-ddCPF1-anti-lacI-lacZ Lac1O-Lac1O; *km*$^R$ in WT-lacZ |
| LacZ-ddCpf1-lacI-Lac1O-Trc2O | pSEL-ddCPF1-anti-lacI-lacZ Lac1O-Trc2O; *km*$^R$ in WT-lacZ |
| LacZ-ddCpf1- | pSEL-ddCPF1-anti-lacI-lacZ J232O-Trc1O; *km*$^R$ in WT-lacZ |

| | |
|---|---|
| lacI-J232O-Trc1O | |
| LacZ-ddCpf1-lacI-J232O-Lac1O | pSEL-ddCPF1-anti-lacI-lacZ J232O-Lac1O; *km*$^R$ in WT-lacZ |
| LacZ-ddCpf1-lacI-J232O-Trc2O | pSEL-ddCPF1-anti-lacI-lacZ J232O-Trc2O; *km*$^R$ in WT-lacZ |
| LacZ-ddCpf1-lacI-J231O-Trc1O | pSEL-ddCPF1-anti-lacI-lacZ J231O-Trc1O; *km*$^R$ in WT-lacZ |
| LacZ-ddCpf1-lacI-J231O-Lac1O | pSEL-ddCPF1-anti-lacI-lacZ J231O-Lac1O; *km*$^R$ in WT-lacZ |
| LacZ-ddCpf1-lacI-Trc1O-tho-Trc1O | pSEL-ddCPF1-anti-lacI-lacZ Trc1O-tho-Trc1O; *km*$^R$ in WT-lacZ |
| LacZ-ddCpf1-lacI-J231O-tho-Trc1O | pSEL-ddCPF1-anti-lacI-lacZ J231O-tho-Trc1O; *km*$^R$ in WT-lacZ |
| Gly-Strain21 | pSEL-ddcpf1-anti-lacI-pslY1-gdh Trc1O-tho-Trc1O; *km*$^R$ in WT |
| Gly-Strain21.1 | NSI::pSI-tho-gly; *spe*$^R$ in Gly-Strain-21 |
| Gly-Strain22 | NSIII::pSIII-glgc-glpf; *cm*$^R$ in Gly-Strain-21.1 |
| HPA-Strain1 | NSIII::pSIII-yqhD; *km*$^R$ in WT |
| HPA-Strain2 | NSIII::pSIII-dhaT; *km*$^R$ in WT |
| HPA-Strain3 | NSIII::pSIII-aox; *km*$^R$ in WT |
| HPA-Strain4 | NSIII::pSIII-alod; *km*$^R$ in WT |
| HPA-Strain5 | NSIII::pSIII-kgsadH-ori; *km*$^R$ in WT |
| HPA-Strain6 | NSIII::pSIII-kgsadH-m; *km*$^R$ in WT |
| HPA-Strain7 | NSIII::pSIII-aldH; *km*$^R$ in WT |
| HPA-Strain8 | NSIII::pSIII-gabD4-ori; *km*$^R$ in WT |
| HPA-Strain9 | NSIII::pSIII-gabD4-m; *km*$^R$ in WT |
| PDO-Strain1 | NSIII::pSIII-trc-DhaB-yqhD; *km*$^R$ in WT |
| PDO-Strain2 | NSI::pSI-psba1-DhaCEFG-96H; *spe*$^R$ in PDO-Strain1 |
| PDO-Strain3 | NSI::pSI-psba1-DhaCEFG-96Q; *spe*$^R$ in PDO-Strain1 |
| PDO-Strain4 | pSEL-cpcB-glpF; *em*$^R$ in PDO-Strain3 |
| PDO-Strain5 | NSII::pSII-DhaBCEFG; *cm*$^R$ in PDO-Strain4 |
| PDO-Strain6 | pSES-ptRNA-Flv3; *cm*$^R$ in PDO-Strain4 |
| PDO-Strain7 | pSES-ptRNA-NdhB; *cm*$^R$ in PDO-Strain4 |
| PDO-Strain8 | pSES-ptRNA-NdhF; *cm*$^R$ in PDO-Strain4 |

| | |
|---|---|
| PDO-Strain9 | NSII::pSII-cpcB-pntAB-eco; $cm^R$ in PDO-Strain4 |
| PDO-Strain10 | NSII::pSII-cpcB-sthA; $cm^R$ in PDO-Strain4 |
| PDO-Strain11 | NSII::pSII-cpcB-glgC; $cm^R$ in PDO-Strain4 |
| 3HP-Strain1 | NSI::pSI-psba1-DhaCEFG-96Q; $spe^R$ in WT |
| 3HP-Strain2 | NSIII::pSIII-trc-DhaB-GabD4; $km^R$ in 3HP-Strain1 |
| 3HP-Strain3 | NSIII::pSIII-tho-DhaB-GabD4; $km^R$ in 3HP-Strain1 |
| 3HP-Strain4 | NSII::pSII-cpcB-ndh2-glpF; $cm^R$ in 3HP-Strain2 |
| 3HP-Strain5 | NSII::pSII-cpcB-Flv1,3-syn; $cm^R$ in 3HP-Strain2 |
| 3HP-Strain6 | NSII::pSII-cpcB-Flv1,3-syu; $cm^R$ in 3HP-Strain2 |
| 3HP-Strain7 | NSII::pSII-cpcB-Flv2,4-syn; $cm^R$ in 3HP-Strain2 |
| 3HP-Strain8 | NSII::pSII-cpcB-pntAB-eco; $cm^R$ in 3HP-Strain2 |
| 3HP-Strain9 | NSII::pSII-cpcB-pntAB-syu; $cm^R$ in 3HP-Strain2 |
| 3HP-Strain10 | NSII::pSII-cpcB-sthA; $cm^R$ in 3HP-Strain2 |
| 3HP-Strain11 | NSII::pSII-cpcB-nox; $cm^R$ in 3HP-Strain2 |
| 3HP-Strain12 | NSII::pSII-dhaT; $cm^R$ in 3HP-Strain2 |
| 3HP-Strain13 | NSII::pSII-yqhD; $cm^R$ in 3HP-Strain2 |
| 3HP-Strain14 | NSII::pSII-cpcB-ndh2-syn; $cm^R$ in 3HP-Strain2 |
| 3HP-Strain15 | NSII::pSII-cpcB-ndh2-bsu; $cm^R$ in 3HP-Strain2 |
| 3HP-Strain16 | pSEL-cpcB-glpF; $em^R$ in 3HP-Strain5 |
| 3HP-Strain17 | NSII::pSII-cpcB-Flv1,3-syn;NSIII::pSIII-lac-DhaB-GabD4;pSEL-cpcB-glpF; $cm^R$, $km^R$ , $em^R$, in 3HP-Strain1 |
| 3HP-Strain18 | NSII::pSII-cpcB-Flv1,3-syn;NSIII::pSIII-cpcB-DhaB-GabD4;pSEL-cpcB-glpF; $cm^R$, $km^R$ , $em^R$, in 3HP-Strain1 |
| 3HP-Strain19 | pSEL-cpc560-Ldh-glpF; $em^R$ in 3HP-Strain5 |
| 3HP-Strain20 | pSEL-cpc560-sAdh-glpF; $em^R$ in 3HP-Strain5 |
| 3HP-Strain21 | pSEL-cpc560-LkLdh-glpF; $em^R$ in 3HP-Strain5 |
| DHA-Strain5 | NSIII::pSIII-gcy1; $km^R$ in WT |
| DHA-Strain1 | NSIII::pSIII-gldA-syu; $km^R$ in WT |
| DHA-Strain3 | NSIII::pSIII-gldA-kpn; $km^R$ in WT |
| DHA-Strain2 | NSIII::pSIII-gldA-eco; $km^R$ inWT |
| DHA-Strain4 | NSIII::pSIII-dhaD-kpn; $km^R$ in WT |
| DHA-Strain6 | NSIII::pSIII-sldAB; $km^R$ in WT |
| DHA-Strain7 | NSIII::pSIII-kpgldA-ecgldA; $km^R$ in WT |
| DHA-Strain8 | pSEL-cpcB-glpF; $em^R$ in WT |
| DHA-Strain9 | NSII::pSII-cpcB-Flv1,3-syn; $cm^R$ in DHA-Strain8 |
| DHA-Strain10 | NSII::pSII-cpcB-Flv1,3-syu; $cm^R$ in DHA-Strain8 |
| DHA-Strain11 | NSII::pSII-cpcB-Flv2,4-syn; $cm^R$ in 3HP-Strain2 |
| DHA-Strain12 | NSII::pSII-cpcB-pntAB-eco; $cm^R$ in DHA-Strain8 |

| | |
|---|---|
| DHA-Strain13 | NSII::pSII-cpcB-pntAB-syu; *cm*$^R$ in DHA-Strain8 |
| DHA-Strain14 | NSII::pSII-cpcB-sthA; *cm*$^R$ in DHA-Strain8 |
| DHA-Strain15 | NSII::pSII-cpcB-nox; *cm*$^R$ in DHA-Strain8 |
| DHA-Strain16 | NSII::pSII-cpcB-ndh2-syn; *cm*$^R$ in DHA-Strain8 |
| DHA-Strain17 | NSII::pSII-cpcB-ndh2-bsu; *cm*$^R$ in DHA-Strain8 |
| Gly-Strain21.2 | NSII::pSII-tho-gly; *spe*$^R$ in Gly-Strain-21 |
| PDO-Strain12 | NSI::pSI-Psba1-DhaCEFG-erm; NSIII::pSIII-trcm-DhaB-yqhD-glgC; *em*$^R$, *cm*$^R$ in Gly-Stran21.2 |
| 3HP-Strain22 | NSI::pSI-Psba1-DhaCEFG-erm; NSIII::pSIII-trcm-haB-GabD4-glgC; *em*$^R$, *cm*$^R$ in Gly-Strain21.2 |
| DHA-Strain18 | NSIII::pSIII-kpgldA-ecgldA-glgC; *cm*$^R$ in Gly-Strain21.2 |

**Table S2. Primers used in this study.**

| SEQUENCE NO. | FORWARD PRIMER | SEQUNENCE FROM 5' TO 3' | REVERSE PRIMER | SEQUNENCE FROM 5' TO 3' | pSII-yqhD | ASSEMBLY METHOD | PLASMID NAME |
|---|---|---|---|---|---|---|---|
| 1 | amp-sept2-R | AAACAAATAGGGGTTCCGCGctagttcggttgaaaggggt | sept2-F | GAATTCCTTAGTTATTCCTATTCTG | pBR-pil-sepT2 | DNA seamless cloning | pBR322m2 |
| 2 | amp-F | CGCGGAACCCCTATTTGTTT | spet2-gj-F | TAGGAATAACTAAGGAATTCCACATTTCCCCGAAAAGTGC | pBR322m | | |
| 3 | nsi-pbr-F | ctaagctgccagccccggcgGAATTCCTTAGTTATTCCTATTC | pbr-gj-R | ATGGAAGCCGGCGGCACCTC | pBR322m2 | DNA seamless cloning | pSIm-cpc560-lacZ |
| 4 | nsi-d-R | cgccggggctggcagcttag | pbr-nsi-F | GAGGTGCCGCCGGCTTCCATatccggcagccggcggagcg | pSI-cpc560-lacZ | | |
| 5 | nsi-pbr-F | ctaagctgccagccccggcgGAATTCCTTAGTTATTCCTATTC | pbr-gj-R | ATGGAAGCCGGCGGCACCTC | pBR322m2 | DNA seamless cloning | pSIm-psba1-lacZ |
| 6 | nsi-d-R | cgccggggctggcagcttag | pbr-nsi-F | GAGGTGCCGCCGGCTTCCATatccggcagccggcggagcg | pSI-psba1-lacZ | | |
| 7 | nsii-F | ggatccacgcattttaattc | nsii-d-R | agcttgtcatctgccggatg | pSII-trc-pilN | DNA seamless cloning | pSIIm-trc-pilN |
| 8 | NSII-gj-F | catccggcagatgacaagctGAATTCCTTAGTTA | nsii-gj-R | gaattaaaatgcgtggatccATGGAAGCCGGCGGCA | pBR322m2 | | |

| | | | | | | | |
|---|---|---|---|---|---|---|---|
| | | TTCCTATTC | | CC | | | |
| 9 | nsiii-R | atcacagtcggcgtcacggc | nsiii-F | tcgaaacccaagccaccctc | pSIII-tps21 | DNA seamless cloning | pSIIIm-tps21 |
| 10 | nsiii-gj-F | gccgtgacgccgactgtgat GAATTCCTTAGTTA TTCCTATTCTGC | nsiii-gj-R | gagggtggcttgggtttcgaCC ACCTCGACCTGAAT GGAA | pBR322m2 | | |
| 11 | cpc560-R | TGAATTAATCTCCT ACTTGACTTTATGA GT | rbcl-F | ACCGGTGTTTGGATT GTCGG | pSIm-cpc560-lacZ | DNA seamless cloning | pSI-cpc560-scegpp2 |
| 12 | cpc560-gpp2-F | TCAAGTAGGAGAT TAATTCAATGGGAT TGACTACTAAACCT | rbcl-gpp2-R | CCGACAATCCAAAC ACCGGTTTACCATTT CAACAGATCGTCC | Saccharomyces cerevisiae S288C | | |
| 13 | cpc560-R | TGAATTAATCTCCT ACTTGACTTTATGA GT | rbcl-F | ACCGGTGTTTGGATT GTCGG | pSIm-cpc560-lacZ | DNA seamless cloning | pSI-cpc560-cregpd2 |
| 14 | cpc560-gpp2-F | TCAAGTAGGAGAT TAATTCAATGATGC TGTCGGGCcgc | rbcl-gpd2-R | CCGACAATCCAAAC ACCGGTCTACACCG AGTTGGACGCGGCG | cregpd2 | | |
| 15 | spe-F | GGATCCCGCACAC CGTGGAA | cpc560-F | ACCTGTAGAGAAGA GTCCCT | pSI-cpc560-scegpp2 | DNA seamless cloning | pSI-cpc560-scegpd1-scegpp2 |
| 16 | spe-cpc560-F | TTCCACGGTGTGC GGGATCCACCTGT AGAGAAGAGTCCC T | gpd1-cpc560-R | CTATCAGCAGCAGC AGACATTGAATTAAT CTCCTACTTGACTTT ATG | pSI-cpc560-scegpp2 | | |
| 17 | cpc-gpd1-R | AGGGACTCTTCTC TACAGGTCTAATCT | gpd1-F | ATGTCTGCTGCTGCT GATAG | Saccharomyces | | |

| | | | | | | | |
|---|---|---|---|---|---|---|---|
| | | TCATGTAGATCTAATTCTT | | | cerevisiae S288C | | |
| 18 | cz-fu-R | AGTCAATCCCCATATAACTACAATCATGTCCGGCAGGTTC | cz-fu-F | TAGTTATATGGGGATTGACTACTAAACCTCTATC | pSI-cpc560-scegpd1-scegpp2 | DNA seamless cloning | pSI-cpc560-Fu-scegpd1-scegpp2 |
| 19 | spe-F | GGATCCCGCACACCGTGGAA | cpc560-F | ACCTGTAGAGAAGAGTCCCT | pSI-cpc560-scegpp2 | DNA seamless cloning | pSI-cpc560-ecogpsA-scegpp2 |
| 20 | spe-cpc560-F | TTCCACGGTGTGCGGGATCCACCTGTAGAGAAGAGTCCCT | syugpsa-cpc560-R | gaagcattacgttggttcatTGAATTAATCTCCTACTTGACTTTATG | pSI-cpc560-scegpp2 | | |
| 21 | cpc560-syugpsA-R | AGGGACTCTTCTCTACAGGTttagtggctgctgcgctc | syugpsa-F | atgaaccaacgtaatgcttc | E. coli MG1655 | | |
| 22 | spe-F | GGATCCCGCACACCGTGGAA | cpc560-F | ACCTGTAGAGAAGAGTCCCT | pSI-cpc560-scegpp2 | DNA seamless cloning | pSI-cpc560-syugpsA-scegpp2 |
| 23 | spe-cpc560-F | TTCCACGGTGTGCGGGATCCACCTGTAGAGAAGAGTCCCT | syugpsa-cpc560-R | gcaacttttgcctcttgcatTGAATTAATCTCCTACTTGACTTTATG | pSI-cpc560-scegpp2 | | |
| 24 | cpc560-syugpsA-R | AGGGACTCTTCTCTACAGGTtcagaccaattccgcttt | syugpsa-F | atgcaagaggcaaaagttgc | Synechococcus elongatus UTEX 2973 | | |
| 25 | spe-F | GGATCCCGCACACCGTGGAA | cpc560-F | ACCTGTAGAGAAGAGTCCCT | pSI-cpc560-scegpp2 | DNA seamless cloning | pSI-cpc560-bsugpsA-scegpp2 |
| 26 | spe-cpc560-F | TTCCACGGTGTGC | bsugpsa- | agcattgtgactttttcatTGA | pSI-cpc560- | | |

| | | | | | | | |
|---|---|---|---|---|---|---|---|
| | | GGGATCCACCTGTAGAGAAGAGTCCCT | cpc560-R | ATTAATCTCCTACTTGACTTTATG | scegpp2 | | |
| 27 | cpc560-bsugpsA-R | AGGGACTCTTCTCTACAGGtttacttcacttgattttcaaacg | bsugpsa-F | atgaaaaaagtcacaatgctt | Bacillus subtilis str. 168 | | |
| 28 | gpsa-cpc560-F | ttgaaaatcaagtgaagtaaACCTGTAGAGAAGAGTCCCTG | gpp2-cpc560-R | GGTTTAGTAGTCAATCCCATTGAATTAATCTCCTACTTGACTTTATG | pSI-cpc560-scegpp2 | DNA seamless cloning | pSI-cpc560-scegpd1-bsugpsA-scegpp2 |
| 29 | trc-gpd1-R | ACACATTATACGAGCCGGATGATTAATTGTCAACAGCTCATCTAATCTTCATGTAGATCTAATTCTTC | gpp2-F | ATGGGATTGACTACTAAACCT | pSI-cpc560-scegpd1-scegpp2 | | |
| 30 | trc-gpsa-F | CCGGCTCGTATAATGTGTGGAATTGTGAGCGGATAACAATTTCACACAatgaaaaaagtcacaatgcttg | gpsa-R | ttacttcacttgattttcaaacgtatt | pSI-cpc560-bsugpsA-scegpp2 | | |
| 31 | gly-cm-R | GGATCCCGCACACCGTGGAAATCGGATCCTACCTGTGACGG | rbcl-RF | TGATGTTCAACTTCGACAGC | pSIIm-trc-pilN | DNA seamless cloning | pSII-cpc560-scegpd1-bsugpsA-scegpp2 |
| 32 | gly-F | TTTCCACGGTGTGCGGGATCC | rbcl-R | GCTGTCGAAGTTGAACATCA | pSI-cpc560-bsugpsA- | | |

| | | | | | | | |
|---|---|---|---|---|---|---|---|
| | | | | | scegpp2 | | |
| 33 | gly-km-R | GGATCCCGCACACCGTGGAAActgatccttcaactcagcaaaag | rbcl-RF | TGATGTTCAACTTCGACAGC | pSIIIm-tps21 | DNA seamless cloning | pSIII-cpc560-scegpd1-bsugpsA-scegpp2 |
| 34 | gly-F | TTTCCACGGTGTGCGGGATCC | rbcl-R | GCTGTCGAAGTTGAACATCA | pSI-cpc560-bsugpsA-scegpp2 | | |
| 35 | gly-F | TTTCCACGGTGTGCGGGATCC | rbcl-R | GCTGTCGAAGTTGAACATCA | pSI-cpc560-bsugpsA-scegpp2 | DNA seamless cloning | pSEL-cpc560-scegpd1-bsugpsA-scegpp2 |
| 36 | rbcl-panl-F | TGATGTTCAACTTCGACAGCccatgatcgagcgatcggcc | erm-amp-F | GCGTTTTTCTTTGTGAGTCCACGCGGAACCCCTATTTGTTT | pSEL-lacZ | | |
| 37 | gly-erm-F | GATCCCGCACACCGTGGAAACGGATGAAGGCACGAACCC | erm-R | TGGACTCACAAAGAAAAAACGC | pBR-pil-sepT2 | | |
| 38 | gg-cm-R | tacggtctccGTAAGAGGTTCCAACTTTCACC | gg-amp-F | tacggtctccgtcccgcggtatcattgcagCAC | pSII-cpc560-scegpd1-bsugpsA-scegpp2 | Golden Gate assembly bsaI | pSII-ptRNA-template |
| 39 | gg-cpcb-F | tacggtctccttacgtgccgatcaCAGTCTCAGCTGCATGCTGgtt | gg-cpcb-F | tacggtctccttacgtgccgatcaCAGTCTCAGCTGCATGCTGgtt | Synechococcus elongatus UTEX 2973 | | |
| 40 | gg-pt1-F | ggctacggtctccgttgaaggaggaattaaccatgcagtggtgGTGGTGGTGGTG | gg-tcr-R | ggctacggtctcgtcaggtcgaggtggcccg | pBR322m2 | | |

| | | | | | | | |
|---|---|---|---|---|---|---|---|
| | | CTGAGACGCACATTTCCCCGAAAAGTGCC | | | | | |
| 41 | gg-rpsl-F | ggctacggtctcgctgagtagccttaagccttaggacgcttc | gg-pt2-R | ggctacggtctctaggaggaattaaccatgcAGTGGTGGTGGTGGTGGTCGAGACGgtagcctcgttgtatatttcttgacacc | E. coli MG1655 | | |
| 42 | gg-rbcl-F | ggctacggtctcttcctaccggtgtttggattgtcggagttgtactcgtcc | gg-amp-R | ggctacggtctccggacccacgctcaccggctcca | pSII-cpc560-scegpd1-bsugpsA-scegpp2 | | |
| 43 | gg-pt-lacZ-F | **TACCGTCTCGGTGC**tggcgaaagggggatgtgct | gg-pt-lacZ-R | **TACCGTCTCATGGTGC**atgattacggattcactggc | pSIm-cpc560-lacZ | Golden Gate assembly bsmbi | pSII-ptRNA-lacZ |
| 44 | cpcB-R | Tcaaccagtctcctgttctc | rbcl-F | ACCGGTGTTTGGATTGTCGG | pSII-ptRNA-template | DNA seamless cloning | pSII-cpcB-pntAB-syu |
| 45 | rbcl-pnt-R | CCGACAATCCAAACACCGGTctaagacaactctttgacagct | cpcB-pnt-F | gagaacaggagactggttgAatgcgatcaccggtcaactg | Synechococcus elongatus UTEX 2973 | | |
| 46 | cpcB-pntAB-eco-F | gagaacaggagactggttgAatgcgaattggcataccaagag | rbcl-pntAB-eco-R | CCGACAATCCAAACACCGGTttacagagctttcaggattgca | E. coli MG1655 | DNA seamless cloning | pSII-cpcB-pntAB-eco |
| 47 | cpcB-R | Tcaaccagtctcctgttctc | rbcl-F | ACCGGTGTTTGGATTGTCGG | pSII-ptRNA-template | | |

| | | | | | | | |
|---|---|---|---|---|---|---|---|
| 48 | cpcB-stha-F | gagaacaggagactggttgAatgccacattcctacgattac | rbcl-stha-R | CCGACAATCCAAACACCGGTttaaaacaggcggtttaaaccgt | E. coli MG1655 | DNA seamless cloning | pSII-cpcB-sthA |
| 49 | cpcB-R | Tcaaccagtctcctgttctc | rbcl-F | ACCGGTGTTTGGATTGTCGG | pSII-ptRNA-template | | |
| 50 | cpcB-glgc-F | GAGAACAGGAGACTGGTTGAATGAAAAACGTGCTGGCGAT | rbcl-glgC-R | CCGACAATCCAAACACCGGTttagatcaccgtgttgtcgg | Synechococcus elongatus UTEX 2973 | DNA seamless cloning | pSII-cpcB-glgC |
| 51 | cpcB-R | Tcaaccagtctcctgttctc | rbcl-F | ACCGGTGTTTGGATTGTCGG | pSII-ptRNA-template | | |
| 52 | cpcB-tpi-F | gagaacaggagactggttgAAtgcgtcggatcatcattgc | rbcl-tpi-R | CCGACAATCCAAACACCGGTttagctctggtaattgacgatg | Synechococcus elongatus UTEX 2973 | DNA seamless cloning | pSII-cpcB-tpi |
| 53 | cpcB-R | Tcaaccagtctcctgttctc | rbcl-F | ACCGGTGTTTGGATTGTCGG | pSII-ptRNA-template | | |
| 54 | cpcB-pfkA-F | gagaacaggagactggttgAatgattaagaaaatcggtgtgttg | rbs-pfkA-R | catttcctcctataggctgattaatacagtttttttcgcgc | Synechococcus elongatus UTEX 2973 | DNA seamless cloning | pSII-cpcB-pfkAB |
| 55 | rbs-pfkB-F | tcagcctataggaggaaatgatggtacgtatctatacgttgac | rbcl-pfkB-R | CCGACAATCCAAACACCGGTttagcgggaaaggtaagcgt | Synechococcus elongatus UTEX 2973 | | |
| 56 | cpcB-R | Tcaaccagtctcctgttctc | rbcl-F | ACCGGTGTTTGGATTGTCGG | pSII-ptRNA-template | | |
| 57 | gg-pt-plsY1-F | **TACCGTCTCGGTGC**gaatgtcaatgcccttgagc | gg-pt-plsY1-R | **TACCGTCTCATGGTGC**atgttgctcagtgttgtcgc | Synechococcus elongatus | Golden Gate | pSII-ptRNA-plsY1 |

| | | | | | | | |
|---|---|---|---|---|---|---|---|
| | | | | | UTEX 2973 | assembly bsmbi | |
| 58 | gg-pt-plsY2-F | **TACCGTCTCGGTGC**ccgaccacttgaacaggtgg | gg-pt-plsY2-R | **TACCGTCTCATGGTGC**atgttggaacagcagcagac | Synechococcus elongatus UTEX 2973 | Golden Gate assembly bsmbi | pSII-ptRNA-plsY2 |
| 59 | gg-pt-eno-F | **TACCGTCTCGGTGC**cttcgaggtggacttctgcc | gg-pt-eno-R | **TACCGTCTCATGGTGC**atgccagacgattacgggac | Synechococcus elongatus UTEX 2973 | Golden Gate assembly bsmbi | pSII-ptRNA-eno |
| 60 | gg-pt-GDH1-F | **TACCGTCTCGGTGC**cgagtgctctggcgcgatcg | gg-pt-GDH1-R | **TACCGTCTCATGGTGC**gtgttacgaggctggggcgc | Synechococcus elongatus UTEX 2973 | Golden Gate assembly bsmbi | pSII-ptRNA-GDH1 |
| 61 | gg-pt-GDH2-F | **TACCGTCTCGGTGC**ggatgtagcgctggggcgag | gg-pt-GDH2-R | **TACCGTCTCATGGTGC**atggaagtttcgacgggcta | Synechococcus elongatus UTEX 2973 | Golden Gate assembly bsmbi | pSII-ptRNA-GDH2 |
| 62 | gg-pt-pgi-F | **TACCGTCTCGGTGC**cgtcatcgaatcccatgcgg | gg-pt-pgi-R | **TACCGTCTCATGGTGC**atgaccgcccagcagctctg | Synechococcus elongatus UTEX 2973 | Golden Gate assembly bsmbi | pSII-ptRNA-pgi |
| 63 | amp-cm-R | **AAACAAATAGGGGTTCCGCGTATAAACGCAGAAAGGCCCA** | pSES-rbcl-R | **gctgtcagtgcccagctcggGCTGTCGAAGTTGAACATCAG** | pSII-ptRNA-template | DNA seamless cloning | pSES-ptRNA-template |

| | | | | | | | |
|---|---|---|---|---|---|---|---|
| 64 | amp-F | **CGCGGAACCCCTATTTGTTT** | pSES-F | **ccgagctgggcactgacagc** | pSES-lacZ | | |
| 65 | amp-cm-R | **AAACAAATAGGGGTTCCGCGTATAAACGCAGAAAGGCCCA** | pSES-rbcl-R | **gctgtcagtgcccagctcggGCTGTCGAAGTTGAACATCAG** | pSII-ptRNA-plsY1 | DNA seamless cloning | pSES-ptRNA-plsY1 |
| 66 | amp-F | **CGCGGAACCCCTATTTGTTT** | pSES-F | **ccgagctgggcactgacagc** | pSES-lacZ | | |
| 67 | amp-cm-R | **AAACAAATAGGGGTTCCGCGTATAAACGCAGAAAGGCCCA** | pSES-rbcl-R | **gctgtcagtgcccagctcggGCTGTCGAAGTTGAACATCAG** | pSII-ptRNA-plsY2 | DNA seamless cloning | pSES-ptRNA-plsY2 |
| 68 | amp-F | **CGCGGAACCCCTATTTGTTT** | pSES-F | **ccgagctgggcactgacagc** | pSES-lacZ | | |
| 69 | amp-cm-R | **AAACAAATAGGGGTTCCGCGTATAAACGCAGAAAGGCCCA** | pSES-rbcl-R | **gctgtcagtgcccagctcggGCTGTCGAAGTTGAACATCAG** | pSII-ptRNA-eno | DNA seamless cloning | pSES-ptRNA-eno |
| 70 | amp-F | **CGCGGAACCCCTATTTGTTT** | pSES-F | **ccgagctgggcactgacagc** | pSES-lacZ | | |
| 71 | amp-cm-R | **AAACAAATAGGGGTTCCGCGTATAAACGCAGAAAGGCCCA** | pSES-rbcl-R | **gctgtcagtgcccagctcggGCTGTCGAAGTTGAACATCAG** | pSII-ptRNA-GDH1 | DNA seamless cloning | pSES-ptRNA-GDH1 |
| 72 | amp-F | **CGCGGAACCCCTATTTGTTT** | pSES-F | **ccgagctgggcactgacagc** | pSES-lacZ | | |

| | | | | | | | |
|---|---|---|---|---|---|---|---|
| 73 | amp-cm-R | **AAACAAATAGGGGTTCCGCGTATAAACGCAGAAAGGCCCA** | pSES-rbcl-R | **gctgtcagtgcccagctcggGCTGTCGAAGTTGAACATCAG** | pSII-ptRNA-GDH2 | DNA seamless cloning | pSES-ptRNA-GDH2 |
| 74 | amp-F | **CGCGGAACCCCTATTTGTTT** | pSES-F | **ccgagctgggcactgacagc** | pSES-lacZ | | |
| 75 | amp-cm-R | **AAACAAATAGGGGTTCCGCGTATAAACGCAGAAAGGCCCA** | pSES-rbcl-R | **gctgtcagtgcccagctcggGCTGTCGAAGTTGAACATCAG** | pSII-ptRNA-pgi | DNA seamless cloning | pSES-ptRNA-pgi |
| 76 | amp-F | **CGCGGAACCCCTATTTGTTT** | pSES-F | **ccgagctgggcactgacagc** | pSES-lacZ | | |
| 77 | dcpf1-917-R | atctatacttaatatatgaacatcatttg | tcr-F | GGTATTCGGAATCTTGCACG | pCPF1-pil | DNA seamless cloning | pdCPF1-pilN |
| 78 | dcpf1-917-F | ttcatatattaagtatagatagaggtgaaagacatttagc | tcr-R | CGTGCAAGATTCCGAATACC | pCPF1-pil | | |
| 79 | km-laci-R | atcagTTTCCACGGTGTGCGtcactgcccgctttccagtc | cz-laci-F | ttCCACAcattatacgagccggatgattaattgtcaatccttcatgtcggcctgcaa | pdCPF1-pilN | DNA seamless cloning | pSEL-ddcpf1-anti-lacZ Trc2O-Trc1O |
| 80 | amp-F | CGCGGAACCCCTATTTGTTT | cr-panl-F | tttatgaaggtcattttttccatgatcgagcgatcggcc | pSEL-lacZ | | |
| 81 | cpf1-crRNA-F | tgcagaataggaataactaattgacaattaatcatccggc | cr-R | aaaaaaatgaccttcataaatcgc | crRNA-lacZ | | |
| 82 | cz-trc-cpf1-F | tccggctcgtataatgTGTGGaattgtgagcgctcacaattTCACACAatgtcaatttat | fncpf1-R | ttagttattcctattctgcacgaa | pdCPF1-pilN | | |

| | | | | | | | |
|---|---|---|---|---|---|---|---|
| | | c | | | | | |
| 83 | km-F | CGCACACCGTGGAAActgat | amp-km-R | AAACAAATAGGGGTTCCGCGttagaaaaactcatcgagcatcaaat | pSIII-cpc560-scegpd1-bsugpsA-scegpp2 | | |
| 84 | fncpf1-R | ttagttattcctattctgcacgaa | cr-panl-F | tttatgaaggtcatttttttccatgatcgagcgatcggcc | pSEL-ddcpf1-lacZ | DNA seamless cloning | pSEL-ddcpf1-anti-lacI-lacZ Trc2O-Trc1O |
| 85 | cpf1-crRNA-F | tgcagaataggaataactaattgacaattaatcatccggc | cr-R | aaaaaaatgaccttcataaatcgc | crRNA-lacI-lacZ | | |
| 86 | fncpf1-R | ttagttattcctattctgcacgaa | cr-panl-F | tttatgaaggtcatttttttccatgatcgagcgatcggcc | pSEL-ddcpf1-anti-lacI-lacZ Trc2O-Trc1O | DNA seamless cloning | pSEL-ddcpf1-anti-lacI-lacZ Trc2O-Lac1O |
| 87 | cpf1-lac-crRNA-F | tgcagaataggaataactaatttacactttatgcttccgg | cr-R | aaaaaaatgaccttcataaatcgc | lac-crRNA-lacI-lacZ | | |
| 88 | trc2O-cpf1-R | tcgcaacgacagaccaaaatatcaattgtgagcgctcacaatttagttattcctattctgcacgaa | cr-panl-F | tttatgaaggtcatttttttccatgatcgagcgatcggcc | pSEL-ddcpf1-anti-lacI-lacZ Trc2O-Trc1O | DNA seamless cloning | pSEL-ddcpf1-anti-lacI-lacZ Trc2O-Trc2O |
| 89 | trc2O-F | attttggtctgtcgttgcgatcgcccgttgcaggccgacatgaaggattgacaattaatcatccgg | cr-R | aaaaaaatgaccttcataaatcgc | crRNA-lacI-lacZ | | |

| | | | | | | | |
|---|---|---|---|---|---|---|---|
| | | c | | | | | |
| 90 | cz-panl-in-F | tccacccataactggcagac | cpf1-lac2O-R | tCCACACaacatacgagccggaagcataaagtgtaaatccttcatgtcggcctgcaacg | pSEL-ddcpf1-anti-lacI-lacZ Trc2O-Trc1O/Lac1O/Trc2O | DNA seamless cloning | pSEL-ddcpf1-anti-lacI-lacZ Lac2O-Trc1O/Lac1O/Trc2O |
| 91 | lac-cpf1-F | tttatgcttccggctcgtatgttGTGTGGaattgtgagcgctcacaattTCACACAatgtcaatttatcaag | cz-panl-in-R | gtctgccagttatgggtgga | pSEL-ddcpf1-anti-lacI-lacZ Trc2O-Trc1O/Lac1O/Trc2O | | |
| 92 | cz-panl-in-F | tccacccataactggcagac | lac1O-j23-F | tCCACACaacatacgagccggaagcataaagtgtaaatccttcatgtcggcctgcaacg | pSEL-ddcpf1-anti-lacI-lacZ Trc2O-Trc1O/Lac1O/Trc2O | DNA seamless cloning | pSEL-ddcpf1-anti-lacI-lacZ Lac1O-Trc1O/Lac1O/Trc2O |
| 93 | lac-cpf1-F | tttatgcttccggctcgtatgttGTGTGGaattgtgagcgctcacaattTCACACAatgtcaatttatcaag | cz-panl-in-R | gtctgccagttatgggtgga | pSEL-ddcpf1-anti-lacI-lacZ Trc2O-Trc1O/Lac1O/Trc2O | | |
| 94 | cz-panl-in-F | tccacccataactggcagac | cpf1-lac2O- | tCCACACaacatacgagcc | pSEL- | DNA | pSEL- |

| | | | | | | | |
|---|---|---|---|---|---|---|---|
| | | | R | ggaagcataaagtgtaaatccttc atgtcggcctgcaacg | ddcpf1-anti-lacI-lacZ Trc2O-Trc1O/Lac1 O/Trc2O | seamless cloning | ddcpf1-anti-lacI-lacZ Lac2O-Trc1O/Lac1 O/Trc2O |
| 95 | lac-cpf1-F | tttatgcttccggctcgtatgtt GTGTGGaattgtgagcgc tcacaattTCACACAatg tcaatttatcaag | cz-panl-in-R | gtctgccagttatgggtgga | pSEL-ddcpf1-anti-lacI-lacZ Trc2O-Trc1O/Lac1 O/Trc2O | | |
| 96 | cz-panl-in-F | tccacccataactggcagac | lac1O-j23-F | tCCACACaacatacgagcc ggaagcataaagtgtaaatccttc atgtcggcctgcaacg | pSEL-ddcpf1-anti-lacI-lacZ Trc2O-Trc1O/Lac1 O/Trc2O | DNA seamless cloning | pSEL-ddcpf1-anti-lacI-lacZ Lac1O-Trc1O/Lac1 O/Trc2O |
| 97 | lac-cpf1-F | tttatgcttccggctcgtatgtt GTGTGGaattgtgagcgc tcacaattTCACACAatg tcaatttatcaag | cz-panl-in-R | gtctgccagttatgggtgga | pSEL-ddcpf1-anti-lacI-lacZ Trc2O-Trc1O/Lac1 O/Trc2O | | |
| 98 | cz-panl-in-F | tccacccataactggcagac | cpf1-J232O-R | CAgctagcattatacctaggact gagctagctgtcaatccttcatgtc ggcctgcaacg | pSEL-ddcpf1-anti-lacI-lacZ | DNA seamless cloning | pSEL-ddcpf1-anti-lacI-lacZ |

| | | | | | | | |
|---|---|---|---|---|---|---|---|
| | | | | | Trc2O-Trc1O/Lac1O/Trc2O | | J232O-Trc1O/Lac1O/Trc2O |
| 99 | J23-cpf1-F | ctcagtcctaggtataatgctagcTGTGGaattgtgagcgctcacaattTCACACAatgtcaatttatcaag | cz-panl-in-R | gtctgccagttatgggtgga | pSEL-ddcpf1-anti-lacI-lacZ Trc2O-Trc1O/Lac1O/Trc2O | | |
| 100 | cz-panl-in-F | tccacccataactggcagac | j231O-laci-F | gcattatacctaggactgagctagctgtcaattgacagctagctcagtcctaggtataatgctagcatctatactggaagagagtcaa | pSEL-ddcpf1-anti-lacI-lacZ Trc2O-Trc1O/Lac1O/Trc2O | DNA seamless cloning | pSEL-ddcpf1-anti-lacI-lacZ J231O-Trc1O/Lac1O/Trc2O |
| 101 | J23-cpf1-F | gcattatacctaggactgagctagctgtcaattgacagctagctcagtcctaggtataatgctagcatctatactggaagagagtcaa | cz-panl-in-R | gtctgccagttatgggtgga | pSEL-ddcpf1-anti-lacI-lacZ Trc2O-Trc1O/Lac1O/Trc2O | | |
| 102 | trc-j23-F | GCTggtatcaccggtaccaattgtgagcgctcacaattcattatacgagccggatgattaattgtcaattgacagctagctcagtcct | cz-panl-in-F | tccacccataactggcagac | pSEL-ddcpf1-anti-lacI-lacZ Trc1O-Trc1O | DNA seamless cloning | pSEL-ddcpf1-anti-lacI-lacZ Trc1O-tho-Trc1O |

| | | | | | | | |
|---|---|---|---|---|---|---|---|
| 103 | tho-cpf1-F | ttggtaccggtgataccAGCATCGTCTTGATGCCCTTGGCAGCACCCTGCTAAGGAGGCAACAAGatgtcaatttatcaagaatttg | cz-panl-in-R | gtctgccagttatgggtgga | pSEL-ddcpf1-anti-lacI-lacZ Trc1O-Trc1O | | |
| 104 | j23-j23-F | ccggtaccaattgtgagcgctcacaattgctagcattatacctaggactgagctagctgtcaattgacagctagctcagtcctaggtataatgctagcatctatactggaagagagtc | cz-panl-in-F | tccacccataactggcagac | pSEL-ddcpf1-anti-lacI-lacZ J231O-Trc1O | DNA seamless cloning | pSEL-ddcpf1-anti-lacI-lacZ J231O-tho-Trc1O |
| 105 | tho-cpf1-F | gcgctcacaattggtaccggtgataccAGCATCGTCTTGATGCCCTTGGCAGCACCCTGCTAAGGAGGCAACAAGatgtcaatttatcaagaatttg | cz-panl-in-R | gtctgccagttatgggtgga | pSEL-ddcpf1-anti-lacI-lacZ J231O-Trc1O | | |
| 106 | tho-cpc560-R | caagggcatcaagacgatgctggtatcaccggtaccaattgtgagGACTTTATGAGTTGGGATTTTCTTAAACAC | nsii-spe-F | cttggtagcaaccgcgctgggTATAAACGCAGAAAGGCCC | pSI-cpc560-scegpd1-bsugpsA-scegpp2 | DNA seamless cloning | pSII-tho-gly |
| 107 | nsii-us-R | ccagcgcggttgctaccaag | tho-gpp2-F | acgctcacaattggtaccggtgataccagcatcgtcttgatgcccttggcagcaccctgctaaggaggca | pSII-cpc560-scegpd1-bsugpsA- | | |

| | | | | | | | |
|---|---|---|---|---|---|---|---|
| | | | | acaagATGGGATTGACTACTAAACCT | scegpp2 | | |
| 108 | tho-gpd1-F | gcatcgtcttgatgcccttggcagcaccctgctaaggaggcaacaagATGTCTGCTGCTGCTGATAG | tho-gpd1-R | ACCTAGGACTGAGCTAGCTGTCAAGAGAGCGTTCACCGACAAACAACAGATAAAACGAAAGGCCCAGTCTTTCGACTGAGCCTTTCGTTTTATTTGCTAATCTTCATGTAGATCTAATTCTTC | pSII-cpc560-scegpd1-bsugpsA-scegpp2 | | |
| 109 | tho-gpsA-F | CTAGCTCAGTCCTAGGTATAATGCTAGCctcacaattggtaccggtgataccagcatcgtcttgatgcccttggcagcaccctgctaaggaggcaacaagatgaaaaaagtcacaatgcttg | tho-gpsA-R | accggtaccaattgtgagcgtatgaaattccacacattatacgagccggatgattaattgtcaacagctcatttacttcacttgattttcaaacgtat | pSII-cpc560-scegpd1-bsugpsA-scegpp2 | | |
| 110 | spe-in-R | ATTCCGTGGCGTTATCCAGC | rbcl-F | ACCGGTGTTTGGATTGTCGG | pSI-cpc560-cregpd2 | DNA seamless cloning | pSI-tho-gly |
| 111 | rbcl-FR | CCGACAATCCAAACACCGGT | spe-in-F | GCTGGATAACGCCACGGAAT | pSII-tho-gly | | |
| 112 | cz-cr2057-F | tgcagaataggaataactaattgacaattaatcatccggc | cz-cr2057-R | ggccgatcgctcgatcatggaaaaaaatgaccttcataaatc | crRNA-Array | DNA seamless cloning | pSEL-ddcpf1-anti-lacI-pslY1-gdh Trc1O- |
| 113 | cpf1-R | ttagttattcctattctgcacgaactc | panl-F | ccatgatcgagcgatcggcc | pSEL-ddcpf1-anti- | | |

| | | | | | | | |
|---|---|---|---|---|---|---|---|
| | | | | | lacI-lacZ Trc1O-tho-Trc1O | | tho-Trc1O |
| 114 | cpcB-glpF-F | GAGAACAGGAGACTGGTTGAATGAgtcaaacatcaacctt | rbcl-glpF-R | CCGACAATCCAAACACCGGTttacagcgaagctttttgtt | E. coli MG1655 | DNA seamless cloning | pSII-cpcB-glpF |
| 115 | cpcB-R | Tcaaccagtctcctgttctc | rbcl-F | ACCGGTGTTTGGATTGTCGG | pSII-ptRNA-template | | |
| 116 | psiii-cz-R | gaccgatcaaccagtccctcatc | nsiii-down-F | cgccagacgcgggtgccca | pSIII-cpc560-scegpd1-bsugpsA-scegpp2 | DNA seamless cloning | pSIII-glgc-glpf |
| 117 | psiii-cm-R | gagggactggttgatcggtcACTAGTgtcccgtCGCAGAAAGGCCCACCCGAA | rbs-glgC-R | AATCGTTTTCTCCTCTGTGTTttagatcaccgtgttgtcgg | pSII-cpcB-glgC | | |
| 118 | rbs-glpf-F | ACACAGAGGAGAAAACGATTatgagtcaaacatcaacctt | nsiii-rbcl-R | ggcacccgcgtctggcgGCTGTCGAAGTTGAACATCA | pSII-cpcB-glpF | | |
| 119 | nsiii-km-R | gagggactggttgatcggtcCTCGAGttagaaaaactcatcgagcatc | psba3-km-F | gtatgtctaaggcgtaatgccttatgagctaataataacaaaactttgcgaattctgatccttcaactcagcaaaag | pSIII-cpc560-scegpd1-bsugpsA-scegpp2 | DNA seamless cloning | pSIII-gabD4-ori |
| 120 | nsiii-UR | gaccgatcaaccagtccctc | rbcl-F | ACCGGTGTTTGGATT | pSIII- | | |

| | | | | | | | |
|---|---|---|---|---|---|---|---|
| | | | | GTCGG | ptGDH1-ptGDH2-glgc-glpf | | |
| 121 | psba3-gabd4-F | gcattacgccttagacatacataaaaattcactggactcaaaacatcTACATAAAAGGAGAATAACCTatgtaccaggatctcgccct | rbcl-gabD4-R | CCGACAATCCAAACACCGGTtcaggcctgggtgatgaact | Cupriavidus necator H16 | | |
| 122 | gabd4-m-1R | GTCGAGACCTGGCTGGGCACGccccag | gabd4-m-2F | GCCGCCCAGATGCTGGCGCGCttc | pSIII-gabD4-ori | DNA seamless cloning | pSIII-gabD4-m |
| 123 | gabd4-m-1F | CCCAGCCAGGTCTCGACCTACCtgat | gabd4-m-2R | GCGCGCCAGCATCTGGGCGGCGGGGtcgatgt | pSIII-gabD4-ori | | |
| 124 | rbs-psba3-R | AGGTTATTCTCCTTTTATGTAgatgttt | Rbcl-F | ACCGGTGTTTGGATTGTCGG | pSIII-gabD4-m | DNA seamless cloning | pSIII-yqhD |
| 125 | RBS-yqhD-F | TACATAAAAGGAGAATAACCTatgaacaactttaatctgcacaccc | rbcl-yqhd-R | CCGACAATCCAAACACCGGTttagcgggcggcttcgtata | E. coli MG1655 | | |
| 126 | rbs-psba3-R | AGGTTATTCTCCTTTTATGTAgatgttt | Rbcl-F | ACCGGTGTTTGGATTGTCGG | pSIII-gabD4-m | DNA seamless cloning | pSIII-aldH |
| 127 | RBS-AldH-F | ACATAAAAGGAGAATAACCTATGAATTTTCATCATCTGGCTT | rbcl-aldh-R | CCGACAATCCAAACACCGGTTTTCAGGCCTCCAGGCTTAT | E. coli MG1655 | | |
| 128 | rbs-psba3-R | AGGTTATTCTCCTTTTATGTAgatgttt | Rbcl-F | ACCGGTGTTTGGATTGTCGG | pSIII-gabD4-m | DNA seamless | pSIII-dhaT |

| | | | | | | | |
|---|---|---|---|---|---|---|---|
| 129 | rbs-dhaT-F | ACATAAAAGGAGAATAACCTatgagctatcgtatgtttgatt | rbcl-dhat-R | CCGACAATCCAAACACCGGTtcagaatgcctggcggaaaa | E. coli MG1655 | cloning | |
| 130 | rbs-psba3-R | AGGTTATTCTCCTTTTATGTAgatgttt | Rbcl-F | ACCGGTGTTTGGATTGTCGG | pSIII-gabD4-m | DNA seamless cloning | pSIII-kgsadH-ori |
| 131 | rbs-kgsadH-F | ACATAAAAGGAGAATAACCTATGGCCAATGTCACGTACAC | rbcl-kgsadh-R | CCGACAATCCAAACACCGGTTTAGACGGCCATGACGGTGA | kgsadH | | |
| 132 | rbs-psba3-R | AGGTTATTCTCCTTTTATGTAgatgttt | Rbcl-F | ACCGGTGTTTGGATTGTCGG | pSIII-gabD4-m | DNA seamless cloning | pSIII-alod |
| 133 | RBS-alod-F | ACATAAAAGGAGAATAACCTATGCGCATCGCCTTTATCGG | rbcl-alod-R | CCGACAATCCAAACACCGGTCTAATCTTTTTTGCGGTAGCCTT | *Pseudomonas sp.* AIU 362 | | |
| 134 | rbs-psba3-R | AGGTTATTCTCCTTTTATGTAgatgttt | Rbcl-F | ACCGGTGTTTGGATTGTCGG | pSIII-gabD4-m | DNA seamless cloning | pSIII-aox |
| 135 | RBS-AOX-F | TACATAAAAGGAGAATAACCTatgcgtatcgcattcatcggc | rbcl-AOX-R | CCGACAATCCAAACACCGGTTCAatccttcttgcgataaccct | *Pseudomonas putida* KT2440 | | |
| 136 | kgsadh-m-F1 | TCGAGTGGTTTGCCGATGAtGGTCGCCGGGTGTATG | kgsadh-m-R2 | GGATGACGGGGTGGGCAATTAAATAGCTGCTGATTTC | pSIII-kgsadH-ori | DNA seamless cloning | pSIII-kgsadH-m |
| 137 | kgsadh-m-F2 | AATTGCCCACCCCGTCATCC | kgsadh-m-R1 | aTCATCGGCAAACCACTCGA | pSIII-kgsadH-ori | | |
| 138 | psba1-R | TTTAAAAGTATTGGGAATTCGATATCAAGCTTatcgatcttgaggttgt | rbcl-F | ACCGGTGTTTGGATTGTCGG | pSIm-psba1-lacZ | DNA seamless cloning | pSI-psba1-DhaCEFG-96Q |

| | | | | | | | |
|---|---|---|---|---|---|---|---|
| | | aaa | | | | | |
| 139 | psba1-dhaC-F | GAATTCCCAATACTTTTAAAGGAGGTTAAAAGTGCAACAGACAACCCAAAT | dhac-R | TCACTCCCTTACTAAGTCGA | Klebsiella pneumoniae DSM 2026 | | |
| 140 | rbs-dhaE-F | TCGACTTAGTAAGGGAGTGATTTAAAGGAGGTTAAAAATGAGCGAGAAAACCATG | dhaF-dhaE-R | ATCCCGGCTATTAACGGCATTTTTAACCTCCTTTAAATTAGCTTCCTTTACGCAGC | Klebsiella pneumoniae DSM 2026 | | |
| 141 | dhaF-F | ATGCCGTTAATAGCCGGGAT | dhaF-R | AAGCGGCCGCTCTAGAACTAGTGGATCCCCCGGGCTGCAGGAATTCCCAATACTTTAATTCGCCTGACCGGCC | Klebsiella pneumoniae DSM 2026 | | |
| 142 | dhaG-F | CTAGTTCTAGAGCGGCCGCTTTAAAGGAGGTTAAAAatgtcgctttcaccgccag | rbcl-dhaG-R | CCGACAATCCAAACACCGGTGAGCTCtcagtttctctcacttaacggc | Klebsiella pneumoniae DSM 2026 | | |
| 143 | amp-in-R | CGTGCACCCAACTGATCTTC | dhag-m-F | tcaccgatagcgacgatcacctgcgtacgctcggcgccaa | pSI-psba1-DHACEFG-96Q | DNA seamless cloning | pSI-psba1-DhaCEFG-96H |
| 144 | amp-in-F | GAAGATCAGTTGGGTGCACG | dhag-m-R | gtgatcgtcgctatcggtga | pSI-psba1-DHACEFG-96Q | | |

| | | | | | | | |
|---|---|---|---|---|---|---|---|
| 145 | trc-km-F | AATTCCACACATTATACGAGCCGGATGATTAATTGTCAACAGCTCATctgatccttcaactcagcaaa | T-psba3-F | TCTGTTGTTTGTCGGTGAACGCTCTCTACTAGAGTCACACTGGCTCACCTTCGGGTGGGCCTTTCTGCGaattcgcaaagttttgttatt | pSIII-gabD4-m | DNA seamless cloning | pSIII-trc-DhaB-gabD4 |
| 146 | trc-dhab-F | CTCGTATAATGTGTGGAATTGTGAGCGGATAACAATTTCACACAATGAAAAGATCAAAACGATTTGC | T-dhaB-R | GTTCACCGACAAACAACAGATAAAACGAAAGGCCCAGTCTTTCGACTGAGCCTTTCGTTTTATTTGATGCCTGGTTATTCAATGGTGTCGGGCT | Klebsiella pneumoniae DSM 2026 | | |
| 147 | trc-km-F | AATTCCACACATTATACGAGCCGGATGATTAATTGTCAACAGCTCATctgatccttcaactcagcaaa | T-psba3-F | TCTGTTGTTTGTCGGTGAACGCTCTCTACTAGAGTCACACTGGCTCACCTTCGGGTGGGCCTTTCTGCGaattcgcaaagttttgttatt | pSIII-yqhD | DNA seamless cloning | pSIII-trc-DhaB-yqhD |
| 148 | trc-dhab-F | CTCGTATAATGTGTGGAATTGTGAGCGGATAACAATTTCACACAATGAAAAGATCAAAACGATTTGC | T-dhaB-R | GTTCACCGACAAACAACAGATAAAACGAAAGGCCCAGTCTTTCGACTGAGCCTTTCGTTTTATTTGATGCCTGGTTATTCAATGGTGTCGGGCT | Klebsiella pneumoniae DSM 2026 | | |

| | | | | | | | |
|---|---|---|---|---|---|---|---|
| 149 | tho-trc-R | gggcatcaagacgatgctggtatcaccggtaccaattgtgagcgtatgaaattgttatccgctcac | amp-in-F | GAAGATCAGTTGGGTGCACG | pSIII-trc-DhaB-gabD4 | DNA seamless cloning | pSIII-tho-DhaB-gabD4 |
| 150 | tho-dhaB-F | ccagcatcgtcttgatgcccttggcagcaccctgctaaggaggcaacaagATGAAAAGATCAAAACGATTTGCAG | amp-in-R | CGTGCACCCAACTGATCTTC | pSIII-trc-DhaB-gabD4 | | |
| 151 | tho-trc-R | gggcatcaagacgatgctggtatcaccggtaccaattgtgagcgtatgaaattgttatccgctcac | amp-in-F | GAAGATCAGTTGGGTGCACG | pSIII-trc-DhaB-yqhD | DNA seamless cloning | pSIII-tho-DhaB-yhqD |
| 152 | tho-dhaB-F | ccagcatcgtcttgatgcccttggcagcaccctgctaaggaggcaacaagATGAAAAGATCAAAACGATTTGCAG | amp-in-R | CGTGCACCCAACTGATCTTC | pSIII-trc-DhaB-yqhD | | |
| 153 | cpcB-R | Tcaaccagtctcctgttctcg | rbcl-F | ACCGGTGTTTGGATTGTCGG | pSII-cpcB-tpi | DNA seamless cloning | pSII-cpcB-ndh2-syn |
| 154 | cpcB-ndh1-F | gagaacaggagactggttgAatgaattcccccacctctcc | ndh1-R | ctaggggcgcactagttcct | Synechocystis sp. PCC 6803 | | |
| 155 | ndh1-ndh2-F | aggaactagtgcgcccctagaatttgattacaaaatgtaacaagccc | ndh3-ndh2-R | tttccatcccactcccgaactcaggaaggttcatttttgagcc | Synechocystis sp. PCC 6803 | | |
| 156 | ndh3-F | gttcgggagtgggatggaaa | rbcl-ndh3-R | CCGACAATCCAAACACCGGTctaccgattcatac | Synechocystis sp. PCC | | |

| | | | | | | | |
|---|---|---|---|---|---|---|---|
| | | | | cggggc | 6803 | | |
| 157 | cpcB-ndh-F | gagaacaggagactggttgAatgtcaaaacatattgtcattctag | rbcl-ndh-R | CCGACAATCCAAACACCGGTttagtaagccaggctgaaaag | Bacillus subtilis str. 168 | DNA seamless cloning | pSII-cpcB-ndh2-bsu |
| 158 | cpcB-R | Tcaaccagtctcctgttctc | rbcl-F | ACCGGTGTTTGGATTGTCGG | pSII-ptRNA-template | | |
| 159 | cpcB-R | Tcaaccagtctcctgttctcg | rbcl-F | ACCGGTGTTTGGATTGTCGG | pSII-ptRNA-template | DNA seamless cloning | pSII-cpcB-Flv1,3-syn |
| 160 | cpcB-flv13-syn-F | gagaacaggagactggttgAgtgggaatccatgcaaaactgg | rbs-flv13-syn-R | TCCTAGGACTGGTAATGAAAActaataatgatcgccagatttcc | Synechocystis sp. PCC 6803 | | |
| 161 | rbs-flv13-syn-F | TTTCATTACCAGTCCTAGGAGTACGGCatgttcactacccccct | rbcl-flv13-syn-R | CCGACAATCCAAACACCGGTttagtaataattgccgactttgcg | Synechocystis sp. PCC 6803 | | |
| 162 | cpcB-R | Tcaaccagtctcctgttctcg | rbcl-F | ACCGGTGTTTGGATTGTCGG | pSII-ptRNA-template | DNA seamless cloning | pSII-cpcB-Flv1,3-syu |
| 163 | cpcB-flv13-syu-F | gagaacaggagactggttgAatggtcgccactaccccag | rbcl-flv13-syu-R | CCGACAATCCAAACACCGGTttagtagtagctgccactcttgcg | Synechococcus elongatus UTEX 2973 | | |
| 164 | cpcB-R | Tcaaccagtctcctgttctcg | rbcl-F | ACCGGTGTTTGGATTGTCGG | pSII-ptRNA-template | DNA seamless cloning | pSII-cpcB-Flv2,4-syn |
| 165 | cpcB-flv24-syn-F | gagaacaggagactggttgAatggttaccctaattgattctcc | rbcl-flv24-syn-R | CCGACAATCCAAACACCGGTtcaaattgatggggactccttacc | Synechocystis sp. PCC 6803 | | |
| 166 | cpcB-R | Tcaaccagtctcctgttctcg | rbcl-F | ACCGGTGTTTGGATTGTCGG | pSII-ptRNA-template | DNA seamless | pSII-cpcB-nox |

| | | | | | | | |
|---|---|---|---|---|---|---|---|
| 167 | cpcB-nox-F | gaacaggagactggttgAAtgaaaatcgtagttatcggtac | rbcl-nox-R | CCGACAATCCAAACACCGGTttatttggcattcaaagctg | Lactococcus lactis subsp. cremoris NZ9000 | cloning | |
| 168 | cm-R | TGATCGGCACGTAAGAGGTT | rbcl-F | ACCGGTGTTTGGATTGTCGG | pSII-ptRNA-template | DNA seamless cloning | pSII-dhaT |
| 169 | cm-psba3-F | AACCTCTTACGTGCCGATCAaattcgcaaagtttgttattat | rbcl-FR | CCGACAATCCAAACACCGGT | pSIII-dhaT | | |
| 170 | amp-T-F | AAACAAATAGGGGTTCCGCGTGGACTCACAAAGAAAAAAC | erm-F | CGCACACCGTGGAAACGGAT | pBR-pil-sepT2 | DNA seamless cloning | pSEL-cpcB-glpF |
| 171 | amp-F | CGCGGAACCCCTATTTGTTT | psel-F | ccatgatcgagcgatcggcc | pSEL-lacZ | | |
| 172 | erm-cpcB-F | ATCCGTTTCCACGGTGTGCGcagtctcagctgcatgctgg | pSEL-rbcl-R | ggccgatcgctcgatcatggGCTGTCGAAGTTGAACATCA | pSII-cpcB-glpF | | |
| 173 | lac-km-F | CACCACACaacatacgagccggaagcataaagtgtaaactgatccttcaactcagcaa | lac-dhab-F | ggctcgtatgttGTGTGGTGCTAAGGAGGCAACAAGATGAAAAGATCAAAACGATT | pSIII-trc-DhaB-gabD4 | DNA seamless cloning | pSIII-lac-DhaB-gabD4 |
| 174 | km-cpcB-F | ttgctgagttgaaggatcagcagtctcagctgcatgctgg | dhab-cpcB-R | AATCGTTTTGATCTTTTCATTcaaccagtctcctgttctc | Synechococcus elongatus UTEX 2973 | DNA seamless cloning | pSIII-cpcB-DhaB-gabD4 |
| 175 | dhab-F | ATGAAAAGATCAA | km-F | ctgatccttcaactcagcaa | pSIII-trc- | | |

| | | | | | | | |
|---|---|---|---|---|---|---|---|
| | | AACGATTTGCAG | | | DhaB-gabD4 | | |
| 176 | cm-R | TGATCGGCACGTAAGAGGTT | rbcl-F | ACCGGTGTTTGGATTGTCGG | pSII-ptRNA-template | DNA seamless cloning | pSII-yqhD |
| 177 | cm-psba3-F | AACCTCTTACGTGCCGATCAaattcgcaaagtttgttattat | rbcl-FR | CCGACAATCCAAACACCGGT | pSIII-yqhD | | |
| 178 | erm-pbr-F | GTTTTTCTTTGTGAGTCCAccagcgcggttgctaccaag | erm-psba3-F | ATCCGTTTCCACGGTGTGCGaattcgcaaagttttgttattat | pSII-yqhD | DNA seamless cloning | pSII-yqhD-erm |
| 179 | erm-R | TGGACTCACAAAGAAAAAACGC | erm-F | CGCACACCGTGGAAACGGAT | pBR-pil-sepT2 | | |
| 180 | rbcl-FR | CCGACAATCCAAACACCGGT | T-psba1-F | CGGGTGGGCCTTTCTGCGctggatttagcgtcttctaat | pSI-psba1-DhaCEFG-96Q | DNA seamless cloning | pSII-DhaBCEFG |
| 181 | cat-trc-F | AACCTCTTACGTGCCGATCAATGAGCTGTTGACAATTAATC | T-dhab-R | CGCAGAAAGGCCCACCCG | pSIII-trc-DhaB-gabD4 | | |
| 182 | cat-F | TGATCGGCACGTAAGAGGTT | rbcl-F | ACCGGTGTTTGGATTGTCGG | pSII-cpcB-yqhD | | |
| 183 | gg-pt-NdhB-F | TACCGTCTCCGTGCTCAAATCCGTGACCAGAAC | gg-pt-NdhB-R | TACCGTCTCGTGGTGCATGGACTTTTAACCC | Synechococcus elongatus UTEX 2973 | Golden Gate assembly bsmbi | pSES-ptRNA-NdhB |
| 184 | gg-pt-NdhF-F | TACCGTCTCAGTGCTGGTGGCCTGGTT | gg-pt-NdhF-R | TACCGTCTCTTGGTGCATGGAATTTCTCTA | Synechococcus elongatus | Golden Gate | pSES-ptRNA- |

| | | | | | | | |
|---|---|---|---|---|---|---|---|
| | | AACGG | | CGATTACG | UTEX 2973 | assembly bsmbi | NdhF |
| 185 | gg-pt-Flv3-F | TACCGTCTCGGTGCAACGGCTGCGATCCCAGT | gg-pt-Flv3-R | TACCGTCTCATGGTGCATGGTCGCCACTACCC | Synechococcus elongatus UTEX 2973 | Golden Gate assembly bsmbi | pSES-ptRNA-Flv3 |
| 186 | gg-pt-glgC-F | TACCGTCTCATGGTGCATGAAAAACGTGCTGGCGAT | gg-pt-glgC-R | TACCGTCTCGGTGCTGCCCGCCAGGGGGACC | Synechococcus elongatus UTEX 2973 | Golden Gate assembly bsmbi | pSES-ptRNA-glgC |
| 187 | cpc560-R | TGAATTAATCTCCTACTTGACTTTATGAG | Rbcl-F | ACCGGTGTTTGGATTGTCGG | pSII-cpc560-scegpd1-bsugpsA-scegpp2 | DNA seamless cloning | pSII-cpc560-LDH |
| 188 | cpc-LDH-F | TCAAGTAGGAGATTAATTCAATGGTTATTCTGATCAACTTCAC | rbcl-LDH-R | CCGACAATCCAAACACCGGTTTAGAATTTGTTTTTATTCAGCGTC | LDH | | |
| 189 | cpc560-R | TGAATTAATCTCCTACTTGACTTTATGAG | Rbcl-F | ACCGGTGTTTGGATTGTCGG | pSII-cpc560-scegpd1-bsugpsA-scegpp2 | DNA seamless cloning | pSII-cpc560-sADH |
| 190 | cpc-sADH-F | TCAAGTAGGAGATTAATTCAATGaaaggtttgcaatgctagg | rbcl-sADH-R | CCGACAATCCAAACACCGGTTTAtaatataactactgctttaatt | Clostridium beijerinckii strain NRRL B593 | | |

| | | | | | | | |
|---|---|---|---|---|---|---|---|
| 191 | cpc560-R | TGAATTAATCTCCTACTTGACTTTATGAG | Rbcl-F | ACCGGTGTTTGGATTGTCGG | pSII-cpc560-scegpd1-bsugpsA-scegpp2 | DNA seamless cloning | pSII-cpc560-lkADH |
| 192 | cpc-lkADH-F | TCAAGTAGGAGATTAATTCAATGactgatcgtttaaaaggc | rbcl-lkADH-R | CCGACAATCCAAACACCGGTTTAttgagcagtgtatccaccatcg | Lactobacillus kefiri DSM 20587 | | |
| 193 | erm-cpc560-F | ATCCGTTTCCACGGTGTGCGaCCTGTAGAGAAGAGTCCCTG | cpcB-rbcl-R | ccagcatgcagctgagactgGCTGTCGAAGTTGAACATCA | pSII-cpc560-LDH | DNA seamless cloning | pSEL-cpc560-Ldh-glpF |
| 194 | erm-F | CGCACACCGTGGAAACGGAT | cpcB-F | cagtctcagctgcatgctgg | pSEL-cpcB-glpF | | |
| 195 | erm-cpc560-F | ATCCGTTTCCACGGTGTGCGaCCTGTAGAGAAGAGTCCCTG | cpcB-rbcl-R | ccagcatgcagctgagactgGCTGTCGAAGTTGAACATCA | pSII-cpc560-sADH | DNA seamless cloning | pSEL-cpc560-sAdh-glpF |
| 196 | erm-F | CGCACACCGTGGAAACGGAT | cpcB-F | cagtctcagctgcatgctgg | pSEL-cpcB-glpF | | |
| 197 | erm-cpc560-F | ATCCGTTTCCACGGTGTGCGaCCTGTAGAGAAGAGTCCCTG | cpcB-rbcl-R | ccagcatgcagctgagactgGCTGTCGAAGTTGAACATCA | pSII-cpc560-lkADH | DNA seamless cloning | pSEL-cpc560-lkAdh-glpF |
| 198 | erm-F | CGCACACCGTGGAAACGGAT | cpcB-F | cagtctcagctgcatgctgg | pSEL-cpcB-glpF | | |
| 199 | rbs-psba3-R | AGGTTATTCTCCTTTTATGTAgatg | rbcl-F | ACCGGTGTTTGGATTGTCGG | pSIII-gabD4-m | DNA seamless cloning | pSIII-gcy1 |
| 200 | psba3-RBS-gcy1-F | ACATAAAAGGAGAATAACCTatgcctgctact | rbcl-gcy1-R | CCGACAATCCAAACACCGGTttacttgaatacttc | Saccharomyces | | |

| | | | | | | | |
|---|---|---|---|---|---|---|---|
| | | ttacatgatt | | gaaaggag | cerevisiae S288C | | |
| 201 | rbs-psba3-R | AGGTTATTCTCCTTTTATGTAgatg | rbcl-F | ACCGGTGTTTGGATTGTCGG | pSIII-gabD4-m | DNA seamless cloning | pSIII-gldA-syu |
| 202 | psba3-RBS-gldA1-F | ACATAAAAGGAGAATAACCTgtgttacgaggctggggcgcac | glda1-R | catTTTTAACCTCCTTTgcgtcattcagccaccacactgt | Synechococcus elongatus UTEX 2973 | | |
| 203 | gldA2-F | cgcAAAGGAGGTTAAAAatggaagtttcgacggg ctacct | rbcl-gldA2-R | CCGACAATCCAAACACCGGTttagccctgtagatgtcgataag | Synechococcus elongatus UTEX 2973 | | |
| 204 | rbs-psba3-R | AGGTTATTCTCCTTTTATGTAgatg | rbcl-F | ACCGGTGTTTGGATTGTCGG | pSIII-gabD4-m | DNA seamless cloning | pSIII-gldA-eco |
| 205 | psba3-RBS-gldA-eco-F | ACATAAAAGGAGAATAACCTatggaccgcattattcaatc | rbcl-gldA-eco-R | CCGACAATCCAAACACCGGTttattcccactcttgcagg | E. coli MG1655 | | |
| 206 | rbs-psba3-R | AGGTTATTCTCCTTTTATGTAgatg | rbcl-F | ACCGGTGTTTGGATTGTCGG | pSIII-gabD4-m | DNA seamless cloning | pSIII-gldA-kpn |
| 207 | psba3-RBS-gldA-kpn-F | ACATAAAAGGAGAATAACCTatggatcgcattattcaatc | rbcl-gldA-kpn-R | CCGACAATCCAAACACCGGTttattcccattcctgcagg | Klebsiella pneumoniae DSM 2026 | | |
| 208 | rbs-psba3-R | AGGTTATTCTCCTTTTATGTAgatg | rbcl-F | ACCGGTGTTTGGATTGTCGG | pSIII-gabD4-m | DNA seamless cloning | pSIII-dhaD-kpn |
| 209 | rbs-kpDhaD-F | ACATAAAAGGAGAATAACCTatgctaaaagttattcaatctccagcc | rbcl-dhaD-R | CCGACAATCCAAACACCGGTttaacgcgccagccactgct | Klebsiella pneumoniae DSM 2026 | | |
| 210 | rbs-psba3-R | AGGTTATTCTCCTT | rbcl-F | ACCGGTGTTTGGATT | pSIII- | DNA | pSIII-sldAB |

| | | | | | | | |
|---|---|---|---|---|---|---|---|
| | | TTATGTAgatg | | GTCGG | gabD4-m | seamless cloning | |
| 211 | rbs-slda-F | ACATAAAAGGAGAATAACCTATGCCCAACACCTACGGCTC | rbs-slda-R | CATTTTTAACCTCCTTTAAATTAGACCGTACCGCGGAGCG | Gluconobacter oxydans 621H | | |
| 212 | rbs-sldb-F | TTTAAAGGAGGTTAAAAATGCGCCGGAGCCATCT | rbcl-sldb-R | CCGACAATCCAAACACCGGTTTAGCCTTTGTGATCGGGC | Gluconobacter oxydans 621H | | |
| 213 | rbs-kpglda-R | ccatTTTTAACCTCCTTTAAAttattcccattcctgcagg | rbcl-F | ACCGGTGTTTGGATTGTCGG | pSIII-gldA-kpn | DNA seamless cloning | pSIII-kpnglda-ecoglda |
| 214 | rbs-ecoglda-F | taaTTTAAAGGAGGTTAAAAatggaccgcattattcaatcac | rbcl-gldA-eco-R | CCGACAATCCAAACACCGGTttattcccactcttgcagg | E. coli MG1655 | | |
| 215 | cpcB-gldA-eco-R | ccagcatgcagctgagactgttattcccactcttgcagg | rbcl-F | ACCGGTGTTTGGATTGTCGG | pSIII-kpnglda-ecoglda | DNA seamless cloning | pSIII-kpgldA-ecgldA-glpF |
| 216 | cpcB-F | cagtctcagctgcatgctgg | rbcl-glpF-R | CCGACAATCCAAACACCGGTttacagcgaagctttttgttc | pSII-cpcB-glpF | | |
| 217 | nsiii-R | gaccgatcaaccagtccctc | psba3-F | aattcgcaaagttttgttattattag | pSIII-kpgldA-ecgldA-glpF | DNA seamless cloning | pSIII-kpgldA-ecgldA-glpF-cm |
| 218 | psba3-cm-F | ataacaaaactttgcgaattTGATCGGCACGTAAGAGGTT | nsiii-cm-R | gagggactggttgatcggtcCGCAGAAAGGCCCACCCG | pSII-cpcB-glpF | | |
| 219 | gg-621h-R | tacgctcttcaTGGCAGG | gg-621-F | tacgctcttccttcCTCGGCC | Gluconobact | Golden | pGOX5-spe- |

| | | | | | | | |
|---|---|---|---|---|---|---|---|
| | | TAAGTCCGGTTC | | AATGTCCAAGC | er oxydans 621H | Gate assembly sapI | cm-km-erm-un |
| 220 | gg-erm-F | tacgctcttcaccaccgaaaaatagCGCACACCGTGGAAACGG | gg-erm-R | tacgctcttcaTATTAAATAATTTATAGCTATTGAAAAGAG | pSEL-cpc560-scegpd1-bsugpsA-scegpp2 | | |
| 221 | gg-spe-F | tacgctcttcaataagtaaCGCACACCGTGGAAACGG | gg-spe-R | tacgctcttccgttgaaggatcagTTATTTGCCGACTACCTTGG | pSI-cpc560-scegpd1-bsugpsA-scegpp2 | | |
| 222 | gg-km-F | tacgctcttccAACTCAGCAAAAGTTCGATTTATTC | gg-km-R | tacgctcttctAGAAAAACTCATCGAGCATC | pSIII-cpc560-scegpd1-bsugpsA-scegpp2 | | |
| 223 | gg-cm-F | gctcttcttctaaCCTACCTGTGACGGAAGATC | gg-cm-R | tacgctcttcgCGCCCCGCCCTGCCACTC | pSII-cpc560-scegpd1-bsugpsA-scegpp2 | | |
| 224 | gg-pjet-F | tacgctcttcggcgtaaCGCTTCCTCGCTCACTGAC | gg-pjet-R | tacgctcttccGAATGGAACCTGCTCCAAGT | pJET1.2 | | |
| 225 | pspe-R | GATGTTTAACTTTGTTTTAGGGCGAC | erm-cds-F | atgaacgagaaaaatataaaacacag | pGOX5-spe-cm-km-erm-un | DNA seamless cloning | pGOX5-spe-cm-km-erm |

| | | | | | | | |
|---|---|---|---|---|---|---|---|
| 226 | erm-p0264-F | CTAAAACAAAGTTAAACATCgatgttcctggatctgttgttc | erm-p0264-R | ttttatatttttctcgttcatttcggtctccctcgccgtaa | Gluconobacter oxydans 621H | | |
| 227 | NSiii-cm-R | gagggactggttgatcggtcCTCGAGTATAAACGCAGAAAGGCCCACCCG | trcm-rbcl-F | TTCCACACATTATACGAGCCGGATGATTAATTGTCAACAGCTCATGCTGTCGAAGTTGAACATCA | pSII-cpcB-glgC | DNA seamless cloning | pSIII-trcm-DhaB-gabD4-glgC |
| 228 | nsiii-u-R | gaccgatcaaccagtccctc | trcm-dhaB-F | GCTCGTATAATGTGTGGAATACATAAAAGGAGAATAACCTATGAAAAGATCAAAACGATTTGC | pSIII-trc-DhaB-gabD4 | | |
| 229 | NSiii-cm-R | gagggactggttgatcggtcCTCGAGTATAAACGCAGAAAGGCCCACCCG | trcm-rbcl-F | TTCCACACATTATACGAGCCGGATGATTAATTGTCAACAGCTCATGCTGTCGAAGTTGAACATCA | pSII-cpcB-glgC | DNA seamless cloning | pSIII-trcm-DhaB-yqhD-glgC |
| 230 | nsiii-u-R | gaccgatcaaccagtccctc | trcm-dhaB-F | GCTCGTATAATGTGTGGAATACATAAAAGGAGAATAACCTATGAAAAGATCAAAACGATTTGC | pSIII-trc-DhaB-yqhD | | |
| 231 | glgc-psba3-F | ccgacaacacggtgatctaaaattcgcaaagttttgttat | rbcl-gldA-eco-R | CCGACAATCCAAACACCGGTttattcccactcttgcagg | pSIII-kpnglda-ecoglda | DNA seamless cloning | pSIII-kpgldA-ecgldA-glgC |

| | | | | | | | |
|---|---|---|---|---|---|---|---|
| 232 | glgc-R | ttagatcaccgtgttgtcgg | rbcl-F | ACCGGTGTTTGGATTGTCGG | pSIII-trcm-DhaB-yqhD-glgC | | |
| 233 | nsi-erm-R | gtgacgagcagggactcgagTGGACTCACAAAGAAAAAACG | psba1-erm-R | cgctaaatccaggcggccgcCGGATGAAGGCACGAACCCA | pSEL-cpc560-scegpd1-bsugpsA-scegpp2 | DNA seamless cloning | pSI-Psba1-DhaCEFG-erm |
| 234 | nsi-us-R | ctcgagtccctgctcgtcac | cz-psba1-F | gcggccgcctggatttagcgtcttc | pSI-psba1-DhaCEFG-96Q | | |

**Table S3. Genes used in this study.**

| | |
|---|---|
| cregpd2 | TCAAGTAGGAGATTAATTCAATGATGCTGTCGGGCCGCACCTGCAAC<br>CATGCCTTCAGCACACGACAGATGAGCCACCAGCGCGGTGCCCTAGC<br>GCTTCGGAGCGCCAGAGTCGCCCAGAGGCCGGTGACGTGTCGCCGA<br>GCTCCCTTTGTGCCCAGCGCCGTGTTTCTGCAGAGCGAACCCGCCCA<br>GAAAACCGCTAGCAGCGCAAACAACGGCGATGCCGCTCCTAGCGAA<br>GCCCGCACTGTGCCCTCGGAACGCGCACTGGCCATCTGGCGCAGCG<br>CCGATGCCGTGTGCTTTGATGTGGATTGCACCATCACCATCAACGATG<br>GCCTGGATCTGCTGGCCGAATTTATGGGCGTGAAAGAAGAAGTGGA<br>AGAACTGACCAACAAAGCCATGGATGGCACCATGAGCCTGACCCGC<br>AGCCTGGAAGAACGCCTGAACCTGATCAACTGCAGCCCCGATGACA<br>TCCGCCGCTTTATCAAAGCCTACCCCCCCCAGAGCCGCCTGGCCCCC<br>GGCATCAAAGAACTGATCAAAGCCCTGCAGAAACGCGGCGTGGCCG<br>TGTACCTGATCAGCGGCGGCTTTCGCGAACTGCTGCTGCCCATCGCC<br>GCCCACCTGGGCATCCCCAAAGATCGCGTGTTTGCCAACCGCATGCA<br>CTGGCAGTGGGATGATGAAACCGGCATGCCCACCAAACTGGTGGGC<br>TTTGATACCAGCGAACCCACCGCCCGCAACCAGGGCAAACCCGAAG<br>CCATCGCCCGCATCCGCGAAAACAACCCCTACAACACCGTGGTGATG<br>ATCGGCGATGGCATCACCGATCTGGAAGCCGTGCAGACCAGCGGCG<br>GCGCCGATCTGTTTATCGGCAGCGGCGTGGTGGTGGAACGCGAAGCC<br>GTGGTTGCCGAAGCTGAGTGGTACGTGTACGATTACAAAGCCCTGGT<br>GAGCGCCCTGAGCCGCTACAAAGTGGCAATGGTGGGCAGCGGCGCG<br>TGGGCCTGCGCTGCCGTGCGCATGATCGCCCAGAACACCAGCCAGG<br>ATGATCCCGAAGATGAATTTGATGATGATGTGCGCATGTGGGTGCACC<br>AGGGCGGCGAACTGGTGGATACCATCAACAGCACCCACGAAAACCC<br>CGCCTACTTTCCAGGGATCCCCCTGGGCCCCAACGTGATCGCCACCG<br>GCAACCTGGCCGAAGCCGTAGCGGATGCCGATCTGCTGGTGTTTTGC<br>GCCCCCCACCAGTACATCCGCGGCATCTGCAAACAGCTGATGGGCAA<br>AGTGAAACCCGGCGCCGCGGCAATCTCGCTGACCAAAGGCATGCGC<br>GTGACCCCCGAAGGCCCCGAACTGATCAGCCAGATCGTGCGCCGCA<br>CACTGGGCGTTGACTGTAGCGTGCTGATGGGCGGCAACATCGCCGAA<br>GATGTGGGCCGCGAACAGCTGAGCGAAGCCGTGATCGGCTACTACA<br>ACCTGGAACACGCCCAGCGCTTTAAAAAACTGTTTCAGCGCCCCTAC<br>TTTCGCGTGACCCTGCTGCCCGATCCCGTGGGCGCCGAACTGTGCGG<br>CACCCTGAAAAACATCGTGGCGCTGGGCGTGGGCATGGTGGATGGC<br>CTGGGCATGGGCCCCAACAGCAAAGCCGCCATCATCCGCCAGGGCC<br>TGCTGGAAATGCGCGATTTTTGCCAGGCCCTGTACCCCAGCGTGCGC<br>GATGATACCTTTCTGGAATGTTGCGGCGTGGGCGATCTCGTGGCCAC<br>CTGTATCGGTGGCCGCAACAGACGCGTAGCCGAAGCATGGACAAGA<br>TCGGCGGTGGAAGGAGCCGAAGCCGGGGAAGGCAACGGCGCCGGC<br>CGCAGCTGGGCCGAACTGGAAAAAGAACTGCTGCAGGGCCAGAAA<br>CTGCAGGGCGTGCTGACCAGCAACGAAGTGCAGCAGATCCTGCGCA<br>CCCGCGGCTGGGAAAGCAAATACCCCCTGTTTACCACCATCAACCGC |

| | |
|---|---|
| | ATCGTGAACGGCCACCTGCCCCCCCACCTGGTGGTGGATTACCTGGAAGGCGCCAAAGCCGACATCGCCGTGGATGTGGAGGAAGACATCGTGCCACTGCCCCGTCAGCCGGCCAGCGCGATGGCCCGCCTATTCGGGCAACTGGTGGGAGGCATTACTCAGCAAGGTGGTGCTGCGGCTGGCGCGGCAGCCAGCGCAGCTGCCGGTGCCGCCAGTGGCGCCGCGTCCAACTCGGTGTAGACCGGTGTTTGGATTGTCGG |
| crRNA-lacZ | ttgacaattaatcatccggctcgtataatgaattgtgagcgctcacaattatgccgtacgatgctgatttaggcaaaaacgggtctaagaactttaaataatttctactgttgtagatcaacgtcgtgactgggaaaaccgtctaagaactttaaataatttctactgttgtagattagcgatttatgaaggtcattttttt |
| crRNA-lacI-lacZ | ttgacaattaatcatccggctcgtataatgaattgtgagcgctcacaattatgccgtacgatgctgatttaggcaaaaacgggtctaagaactttaaataatttctactgttgtagatcaacgtcgtgactgggaaaaccgtctaagaactttaaataatttctactgttgtagatTTGATGTCTCTGACCAGACACCgtctaagaactttaaataatttctactgttgtagattagcgatttatgaaggtcattttttt |
| lac-crRNA-lacI-lacZ | tttacactttatgcttccggctcgtatgttGaattgtgagcgctcacaattatgccgtacgatgctgatttaggcaaaaacgggtctaagaactttaaataatttctactgttgtagatcaacgtcgtgactgggaaaaccgtctaagaactttaaataatttctactgttgtagatTTGATGTCTCTGACCAGACACCgtctaagaactttaaataatttctactgttgtagattagcgatttatgaaggtcattttttt |
| crRNA-A | ttgacaattaatcatccggctcgtataatgaattgtgagcgctcacaattatgccgtacgatgctgatttaggcaaaaacgggtctaagaactttaaataatttctactgttgtagatgacgggctaccttccgcaggcagtctaagaactttaaataatttctactgttgtagatacggcgatcgcttcagaacaatgtctaagaactttaaataatttctactgttgtagatggctccttcccagcaggctatcgtctaagaactttaaataatttctactgttgtagatTTGATGTCTCTGACCAGACACCgtctaagaactttaaataatttctactgttgtagattagcgatttatgaaggtcattttttt |

| | |
|---|---|
| rr<br>a<br>y | |
| k<br>g<br>s<br>a<br>d<br>h | ATGGCCAATGTCACGTACACCGACACCCAACTGCTGATTGATGGCGA<br>ATGGGTCGATGCGGCGTCGGGCAAAACCATTGATGTGGTGAACCCCG<br>CCACGGGCAAACCCATTGGTCGCGTGGCGCATGCCGGTATCGCCGAT<br>CTGGATCGCGCTTTAGCCGCCGCGCAATCGGGTTTTGAGGCTTGGCG<br>CAAAGTCCCGGCCCATGAACGCGCGGCCACCATGCGCAAAGCCGCG<br>GCGCTGGTCCGCGAACGGGCCGATGCGATTGCCCAGCTGATGACGC<br>AAGAACAAGGCAAGCCTTTAACCGAAGCCCGCGTGGAGGTTTTAAG<br>CGCGGCCGACATTATCGAGTGGTTTGCCGATGAAGGTCGCCGGGTGT<br>ATGGTCGCATTGTCCCCCCCCGGAATCTGGGCGCGCAACAAACCGTC<br>GTGAAAGAACCGGTGGGCCCCGTCGCCGCCTTTACCCCTTGGAACTT<br>CCCCGTGAATCAAGTGGTGCGCAAACTGTCGGCGGCGCTGGCGACC<br>GGTTGTAGCTTTCTGGTCAAAGCGCCCGAGGAAACCCCCGCGAGCC<br>CGGCCGCTTTACTGCGCGCCTTTGTCGATGCCGGTGTGCCGGCCGGT<br>GTGATCGGTCTGGTCTACGGCGACCCCGCCGAAATCAGCAGCTATTT<br>AATTcCCCACCCCGTCATCCGCAAAGTCACGTTCACCGGCTCGACGC<br>CCGTGGGCAAACAACTGGCCTCTTTAGCCGGTTTACACATGAAACGC<br>GCCACGATGGAACTGGGCGGTCATGCCCCCGTGATCGTGGCCGAAG<br>ATGCGGATGTCGCTTTAGCGGTGAAAGCCGCGGGCGGCGCGAAATTT<br>CGCAATGCCGGCCAAGTTTGCATTAGCCCCACCCGCTTTTTAGTGCA<br>CAACTCGATCCGCGATGAGTTTACCCGCGCTTTAGTCAAACACGCCG<br>AAGGTTTAAAGGTGGGCAACGGTTTAGAAGAAGGCACTACGCTGGG<br>CGCCCTCGCCAACCCCCGCCGTTTAACCGCCATGGCCAGCGTCATCG<br>ATAACGCCCGCAAGGTCGGTGCCAGCATTGAGACCGGCGGCGAACG<br>CATCGGCAGCGAGGGCAACTTTTTCGCCCCCACCGTCATCGCCAATG<br>TCCCGCTGGACGCCGATGTCTTCAACAATGAACCGTTTGGCCCGGTC<br>GCGGCCATTCGCGGCTTTGATAAGCTGGAGGAAGCGATTGCCGAGGC<br>GAATCGTTTACCCTTTGGTCTCGCCGGCTATGCGTTCACGCGCAGCTT<br>TGCCAACGTGCATCTGCTGACCCAACGGCTGGAGGTGGGCATGCTGT<br>GGATCAACCAACCGGCGACGCCTTGGCCCGAAATGCCGTTTGGCGG<br>CGTCAAGGATAGCGGCTATGGCAGCGAAGGCGGTCCGGAGGCTTTA<br>GAACCGTATTTAGTGACCAAGAGCGTCACCGTCATGGCCGTCTAA |
| L<br>D<br>H | ATGGTTATTCTGATCAACTTCACGGAGGTGGGCTTTATGACGAAGGT<br>GTTCGCCTACGCCATCCGCAAGGATGAGGAACCGTTTCTGAATGAGT<br>GGAAGGAAGCGCACAAGGATATCGACGTTGACTACACCGACAAACT<br>GCTGACCCCGGAAACGGCGAAACTGGCGAAAGGTGCCGACGGCGT<br>GGTGGTTTACCAGCAGCTGGACTATACCGCCGATACGCTGCAAGCGC<br>TGGCCGATGCGGGCGTTACGAAGATGAGTCTGCGCAACGTGGGCGT<br>GGACAACATCGACATGGACAAAGCGAAGGAGCTGGGCTTCCAGATT<br>ACCAACGTGCCGGTGTACAGCCCAAATGCCATCGCCGAACATGCGGC<br>GATTCAAGCCGCCCGTGTTCTCCGTCAAGATAAACGCATGGACGAGA<br>AGATGGCGAAGCGCGATCTGCGTTGGGCGCCAACCATTGGTCGTGA |

| | |
|---|---|
| | AGTGCGCGATCAAGTTGTTGGTGTTGTTGGCACCGGTCATATCGGCC<br>AAGTTTTCATGCGCATCATGGAGGGCTTTGGCGCCAAGGTGATCGCC<br>TACGACATCTTCAAGAACCCGGAGCTGGAGAAGAAGGGTTACTACG<br>TGGACAGTCTGGATGATCTGTACAAGCAAGCCGATGTTATCAGTCTG<br>CACGTGCCGGATGTGCCGGCCAACGTGCACATGATCAACGACAAGA<br>GTATCGCGGAGATGAAAGATGGCGTTGTGATCGTGAACTGCAGTCGT<br>GGCCCGCTGGTTGACACCGATGCCGTGATTCGCGGTCTGGACAGCGG<br>CAAAATCTTCGGCTTCGTGATGGACACCTACGAGGACGAAGTGGGC<br>GTTTTCAACAAGGATTGGGAGGGCAAAGAGTTCCCGGACAAACGTC<br>TGGCGGATCTGATCGATCGTCCGAATGTGCTGGTTACCCCGCATACCG<br>CCTTCTACACCACCCATGCCGTTCGCAACATGGTGGTGAAGGCCTTC<br>AACAACAATCTGAAGCTGATCAACGGCGAAAAACCGGACAGCCCGG<br>TGACGCTGAATAAAAACAAATTCTAA |
